# Lysine63-linked ubiquitin chains earmark GPCRs for BBSome-mediated removal from cilia

**DOI:** 10.1101/2020.03.04.977090

**Authors:** Swapnil Rohidas Shinde, Andrew R. Nager, Maxence V. Nachury

## Abstract

Regulated trafficking of G-protein coupled receptors (GPCRs) controls cilium-based signaling pathways. β-arrestin, a molecular sensor of activated GPCRs, and the BBSome, a complex of Bardet-Biedl Syndrome (BBS) proteins, are required for the signal-dependent exit of ciliary GPCRs but the functional interplay between β-arrestin and the BBSome remains elusive. Here we find that, upon activation, ciliary GPCRs become tagged with K63-linked ubiquitin (K63Ub) chains in a β-arrestin-dependent manner prior to BBSome-mediated exit. Removal of ubiquitin acceptor residues from the somatostatin receptor 3 (SSTR3) and from the orphan GPCR GPR161 demonstrates that ubiquitination of ciliary GPCRs is required for their regulated exit from cilia. Furthermore, targeting a K63Ub-specific deubiquitinase to cilia blocks the exit of GPR161, SSTR3 and Smoothened (SMO) from cilia. Finally, ubiquitinated proteins accumulate in cilia of mammalian photoreceptors and *Chlamydomonas* cells when BBSome function is compromised. We conclude that K63Ub chains mark GPCRs and other unwanted ciliary proteins for recognition by the ciliary exit machinery.

## INTRODUCTION

The regulated trafficking of signaling receptors in and out of cilia is a central regulatory mechanism of many cilia-based signaling pathways (Nachury and Mick, 2019; Anvarian et al., 2019; Mykytyn and Askwith, 2017; Gigante and Caspary, 2020). For example, upon activation of the Hedgehog signaling pathway, the Hedgehog receptor Patched 1 and the G protein-coupled receptor (GPCR) GPR161 disappear from cilia while GPCR Smoothened (SMO) accumulates in cilia (Rohatgi et al., 2007; Corbit et al., 2005; Mukhopadhyay et al., 2013). In all three cases, regulated exit from cilia plays a major role in the on-demand redistribution of signaling molecules (Nachury and Mick, 2019). Signal-dependent exit is likely to be a general characteristic of ciliary GPCRs as the Somatostatin Receptor 3 (SSTR3), the Dopamine Receptor 1 (D1R), the Melanocortin concentrating hormone receptor 1 (MCHR1) and the neuropeptide receptor 2 (NPY2R) all disappear from cilia upon exposure to agonist (Domire et al., 2011; Loktev and Jackson, 2013; Nager et al., 2017; Green et al., 2015). A major question is how GPCRs are selected for removal from cilia in an activity-dependent manner.

The major trafficking entities mediating signal-dependent exit from cilia are the BBSome and β-arrestin 2. The BBSome is an evolutionarily conserved complex of eight Bardet-Biedl Syndrome (BBS) proteins that directly recognizes intracellular determinants in GPCRs and ferries them out of cilia (Nachury, 2018; Wingfield et al., 2018). The BBSome associates with the intraflagellar transport (IFT) machinery and has been proposed to act as adaptor between the motor-driven intraflagellar transport trains and the membrane proteins that are to be removed from cilia. Because the BBSome is not known to have the ability to discriminate active from inactive GPCRs, there must exist another layer of regulation that commits activated GPCRs for exit. β-arrestin 2 is a well-established molecular sensor of the activation state of GPCRs that is required for the signal-dependent exit of GPR161 and SSTR3 (Pal et al., 2016; Green et al., 2015; Nager et al., 2017). No association of β-arrestin 2 with ciliary trafficking complexes has been reported to date and it remains unclear how β-arrestin 2 relays information regarding the state of activation of ciliary GPCRs to the ciliary exit machinery.

An emerging player in ciliary exit is ubiquitination (Shearer and Saunders, 2016). Ubiquitin (Ub), a 76-amino acid polypeptide, becomes conjugated to acceptor lysine residues on substrate proteins by ubiquitin ligases and tags substrates for degradation or other regulatory fates (Yau and Rape, 2016; Swatek and Komander, 2016). A role for ubiquitin in promoting ciliary exit is suggested by multiple lines of evidence. First, interfering with Patched1 ubiquitination blocks its signal-dependent exit from cilia (Kim et al., 2015; Yue et al., 2014). Second, fusing ubiquitin to PKD-2, the olfactory receptor ODR-10 or the TRP channel OSM-9 at their cytoplasmic end results in the disappearance of these proteins from cilia (Hu et al., 2007; Xu et al., 2015). The accumulation of these ubiquitin fusions inside cilia when BBSome function is compromised suggests that the BBSome might sort ubiquitinated signaling receptors out of cilia, in line with a reported association of BBSome with ubiquitinated proteins in trypanosomes (Langousis et al., 2016). Interestingly, TRIM32 is a ubiquitin ligase mutated in BBS patients (Chiang et al., 2006). Third, the ubiquitin ligase Cbl is recruited to cilia upon activation of the ciliary tyrosine kinase receptor PDGFRαα and Cbl is required for termination of PDGFRαα signaling (Schmid et al., 2018). Last, a dramatic rise in ubiquitination of the ciliary proteome is observed when *Chlamydomonas* cilia disassemble (Huang et al., 2009). In particular, α-tubulin becomes ubiquitinated on K304 upon cilia disassembly and expression of α-tubulin[K304R] slows down cilia disassembly (Wang et al., 2019).

After conjugation onto substrates, ubiquitin itself often serves as a substrate for ubiquitination. The resulting ubiquitin chains are characterized by the acceptor lysine residue on ubiquitin with lysine 48-linked Ub (UbK48) chains targeting soluble proteins for proteasomal degradation while lysine 63-linked Ub (UbK63) chains assembled onto membrane proteins constitute the major driver for sorting from the limiting membrane of late endosomes into intralumenal vesicles and ultimately lysosomal degradation (Piper et al., 2014; Yau and Rape, 2016; Swatek and Komander, 2016). Intriguingly, in shortening *Chlamydomonas* cilia K63Ub linkages predominate at least 5-fold over K48 linkages and only K63Ub linkages are detected on α-tubulin. A role for K63Ub chains in ciliary trafficking remains to be determined.

Together, these data led us to investigate the interplay between ubiquitination, β-arrestin 2 and the ciliary exit machinery. Using a combination of cellular imaging, biochemical studies and cilia-specific manipulations, we find that β-arrestin 2 directs the activation-dependent addition of K63-linked ubiquitin chains onto ciliary GPCRs that are then selected by the BBSome for removal from cilia.

## RESULTS

### Signal-dependent ubiquitination of ciliary GPCRs is required for their regulated exit from cilia

As a first step in investigating the interplay between ubiquitination and the ciliary exit machinery, we sought to determine whether activation of ciliary GPCRs leads to their ubiquitination inside cilia prior to retrieval into the cell. We blocked BBSome-dependent exit by deleting its cognate GTPase ARL6/BBS3 in mouse inner medulla collecting duct (IMCD3) cells. IMCD3 cells been previously characterized (Liew et al., 2014; Ye et al., 2018). As in previous studies (Ye et al., 2018; Nager et al., 2017), all GPCRs were expressed at near-endogenous levels by driving expression with extremely weak promoters. The fluorescent protein mNeonGreen (NG) fused to the cytoplasm-facing C-terminus of GPCRs allowed direct visualization. A biotinylation Acceptor Peptide (AP) fused to the extracellular N-terminus biotinylated by an ER-localized biotin ligase (BirA-ER) enabled pulse labeling of surface-exposed molecules and denaturing purifications. When SSTR3 was expressed in *Arl6^-/-^* IMCD3 cells, addition of its agonist somatostatin (sst) led to a drastic increase in the ciliary levels of ubiquitin observed upon immunostaining with the well-characterized FK2 (**Figure 1A-B**) and FK1 (**Figure S1A-B**) monoclonal antibodies (Fujimuro et al., 1994; Emmerich and Cohen, 2015; Haglund et al., 2003). All further experiments were conducted with the FK2 antibody. *Arl6^-/-^* cells accumulated Ub signals inside cilia in the absence of SSTR3 expression (**Figure 1A-B**), demonstrating that some endogenous proteins become ubiquitinated inside the cilia of *Arl6^-/-^* IMCD3 cells. No ubiquitin signal was detected inside cilia when BBSome function was intact (**Figure 1A-B and S1A-D**), indicating that the BBSome efficiently removes ubiquitinated proteins from cilia. Because the sst-dependent increase in ciliary ubiquitin signal depends upon SSTR3 expression (**Figure 1A-B**), these results suggest that either SSTR3 itself or a downstream effector of SSTR3 become ubiquitinated inside cilia upon activation.

**Figure 1.**
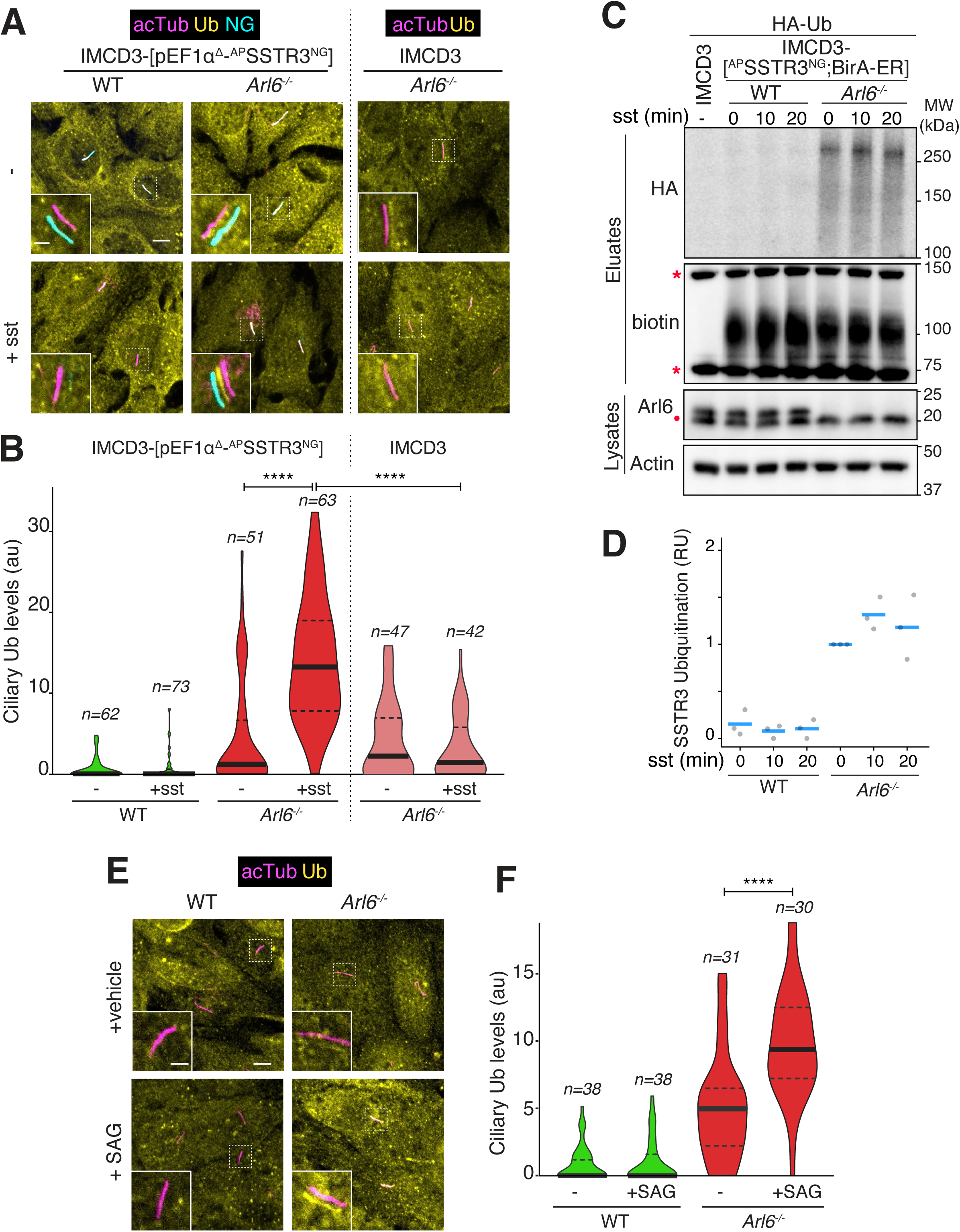
Ubiquitinated signaling receptors accumulate inside cilia of *Arl6^-/-^* cells. **A.** SSTR3 fused to the fluorescent protein mNeonGreen (NG) at its intracellular C-terminus and a biotinylation Acceptor Peptide (AP) tag at its extracellular N-terminus was expressed under the control of an attenuated EF1α promoter (pEF1α^Δ^) in wild type (WT) and *Arl6^-/-^* IMCD3 cells. Cells were treated with or without somatostatin-14 (sst) for 2 h, then fixed and stained for acetylated tubulin (acTub, magenta) and ubiquitin (Ub, yellow). ^AP^SSTR3^NG^ (cyan) was imaged through the intrinsic fluorescence of NG. Channels are shifted in the insets to facilitate visualization of overlapping ciliary signals. Scale bar: 5μm (main panel), 2μm (inset). In WT cells, the ciliary SSTR3 signal decreases over the experimental time course. In *Arl6^-/-^* cells, SSTR3 fails to exit cilia and an increase in the ciliary Ub level is detected. As a control, *Arl6^-/-^* cells that did not express SSTR3-NG were tested; no increase in ciliary Ub levels was observed upon addition of sst. **B.** The fluorescence intensity of the Ub channel in the cilium was measured in each condition and the data are represented as violin plots. The thick bar indicates the median and the dotted lines the first and third quartiles. An 11-fold increase in ciliary Ub signal is observed upon addition of sst to SSTR3-expressing cells. Asterisks indicate ANOVA significance value. ****, p= <0.0001. **C.** WT or *Arl6^-/-^* IMCD3 cells stably expressing ^AP^SSTR3^NG^ and the biotin ligase BirA targeted to the ER lumen (BirA-ER) were transiently transfected with HA-tagged ubiquitin (HA-Ub). 10 µM biotin was added to cells 24 h post transfection for maximal biotinylation of ^AP^SSTR3^NG^ and, after another 18 h, cells were treated with sst (10μM) for indicated times. Cells were lysed under denaturing conditions and biotinylated SSTR3 was captured on streptavidin resin. Eluates were probed for HA via immunoblotting and for biotin via streptavidin-HRP. Two major biotinylated proteins endogenous to cells are marked by asterisks. Whole cell lysates were probed for Arl6 and, as a loading control, actin. A non-specific band cross-reacting with the anti-Arl6 antibody is marked with a dot. **D.** Quantitation of SSTR3 ubiquitination. The signals of HA-Ub conjugated to SSTR3 in the streptavidin eluates were measured. The experiment shown in **C** was repeated three times and for each experiment, Ub-SSTR3 signals were normalized to the value in *Arl6^-/-^* cells at t = 0 of sst stimulation and plotted as grey circles. The horizontal blue lines represent mean values. **E.** IMCD3 cells of the indicated genotypes were treated with the Smoothened agonist SAG or the vehicle DMSO for 2h. Cells were then fixed and stained for acTub and Ub. Channels are shifted in the insets to facilitate visualization of overlapping ciliary signals. Scale bar 5μm (main panel), 2μm (inset). Activation of Hh signaling promotes a detectable increase in ciliary Ub levels only in *Arl6^-/-^* cells. **F.** Violin plots of the fluorescence intensity of the Ub channel in the cilium in each condition are shown. Asterisks indicate ANOVA significance value. ****, p= <0.0001.

To determine whether SSTR3 becomes ubiquitinated in response to sst, we biochemically ubiquitin via an HA tag on transfected HA-Ub. As typical for a glycosylated protein, SSTR3 migrated as a broad band centered around 100 kDa (**Figure 1C**). The amount of SSTR3 recovered did not change appreciably between WT and *Arl6^-/-^* cells or upon stimulation with sst (**Figure 1C**). (**Figure 1C**). Remarkably, while only very faint Ub signals were detected associated with SSTR3 in WT cells, a Ub smear extending from 100 kDa upward was detected in the SSTR3 pull-downs from *Arl6^-/-^* cells. Furthermore, stimulation with sst resulted in a modest but reproducible increase in SSTR3 ubiquitination in *Arl6^-/-^* cells (Figure 1D). We conclude that the BBSome/ARL6 system is required for the degradation of ubiquitinated SSTR3 and that SSTR3 becomes ubiquitinated in response to stimulation with sst.

We next sought to test whether signaling receptors endogenous to IMCD3 cells are ubiquitinated inside cilia in an activity-dependent manner. One signaling pathway that is natively expressed in IMCD3 cells is the Hedgehog pathway. Prior experiments have detected normal trafficking dynamics of SMO and GPR161 in IMCD3 cells (Ye et al., 2018; Mukhopadhyay et al., 2013) and **Figure 4D**). As GPR161 and SMO both accumulate in cilia of *Arl6^-/-^* cells without Hh pathway stimulation (Liew et al., 2014; Zhang et al., 2011), the ubiquitin signal detected in *Arl6^-/-^* cilia may correspond to ubiquitin conjugated to SMO or GPR161. We detected a significant elevation of the ciliary ubiquitin levels in *Arl6^-/-^* cells when treated with the SMO agonist SAG (**Figure 1E-F**). Because SMO ubiquitination was previously shown to decrease upon its activation (Xia et al., 2012; Jiang et al., 2019), it is likely that the elevated ciliary ubiquitin signal detected in SAG-treated cells is associated with GPR161. As activation of SMO triggers exit of GPR161 but this exit is frustrated when BBSome function goes awry, these data suggest that GPR161 becomes ubiquitinated prior to its BBSome-dependent exit. Collectively, these data suggest that GPCRs are ubiquitinated in a regulated fashion inside cilia and subsequently retrieved into the cell by the BBSome.

To test the functional importance of GPCR ubiquitination in BBSome-mediated exit from cilia, we removed all ubiquitination sites from SSTR3 and GPR161 by mutating all cytoplasm-exposed lysine residues to arginine (cK0 variants). Exit kinetics were precisely monitored by real-time tracking of individual cilia in live cells. In both cases, the cK0 variant underwent markedly slower signal-dependent exit from cilia than the WT allele (**Figure 2A-B**). Measurement of exit rates revealed that exit of SSTR3cK0 was ∼2-fold slower than SSTR3 and that exit of GPR161cK0 was ∼2.5-fold slower than GPR161. The residual exit rates of the cK0 mutants indicate the existence of alternative mechanisms of exit that complement GPCR ubiquitination or bypass BBSome-dependent retrieval (see discussion).

**Figure 2.**
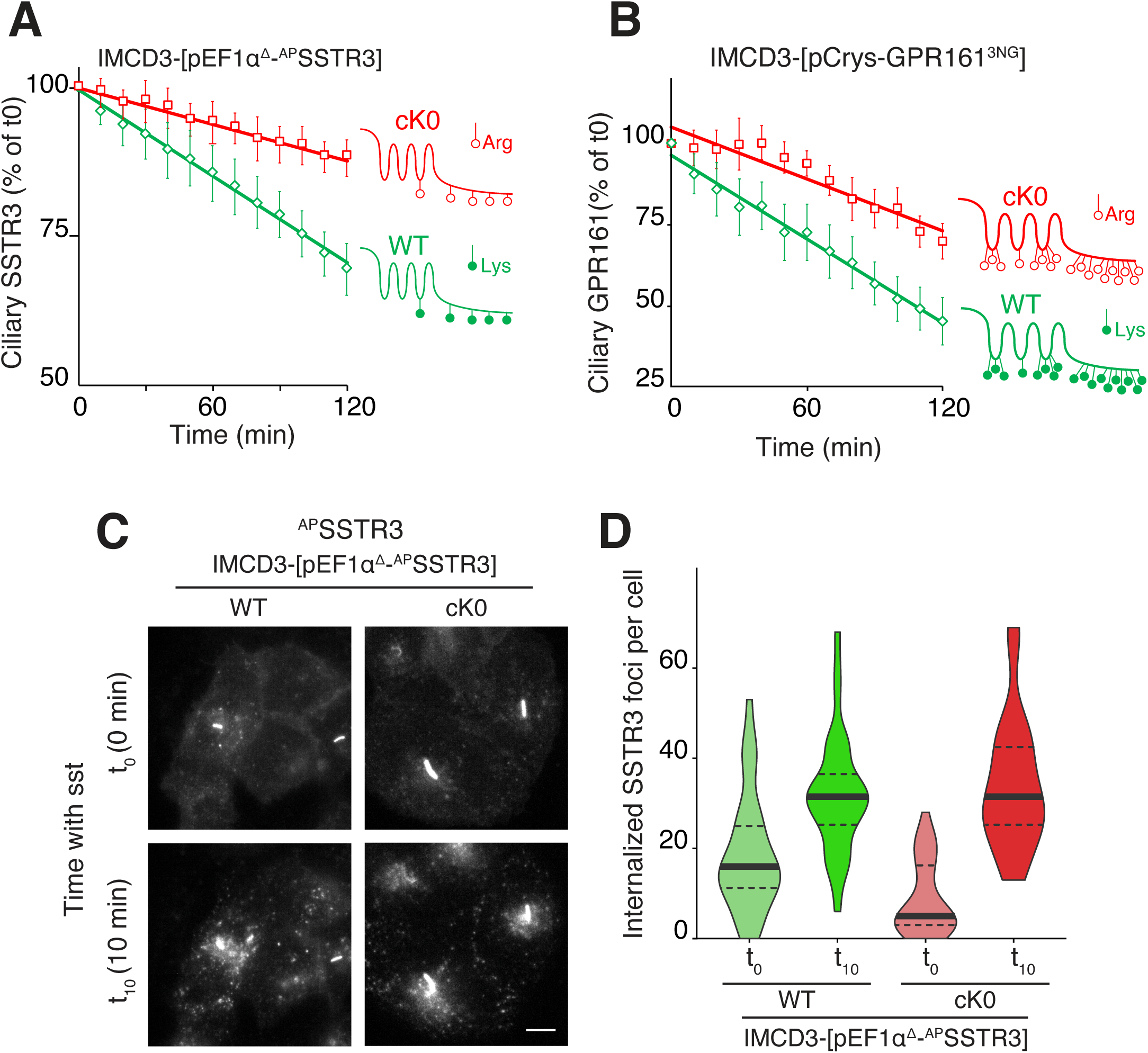
Ubiquitination of ciliary GPCRs is required for signal-dependent exit but not for endocytosis. **A.** IMCD3-[pEF1α^Δ^-^AP^SSTR3] cells stably expressed the ER-localized biotin ligase BirA to enables the biotinlylation of ^AP^SSTR3. SSTR3 was either the wild-type allele (WT) or a variant where all five cytoplasm-facing lysine residues (listed in Methods) were mutated to arginine (cK0). Ciliary ^AP^SSTR3 was pulse-labeled by addition of Alexa647-conjugated mSA (mSA647) to the medium for 5-10 min before addition of sst. Far red fluorescence was tracked in individual cilia at 10 min capture intervals. For each individual cilia, fluorescence intensities were normalized to the value at t = 0. A comparison of the ciliary levels of SSTR3 at t=0 is shown in **Fig. S2A**. Data were plotted and fitted to a line. Error bars: 95% CI. *N* =18-22 cilia. **B.** GPR161 fused to three tandem repeats of NG at its C-terminus was expressed under the control of the ultra-weak δ-crystallin promoter (pCrys) in IMCD3 cells. GPR161 was either the wild-type allele (WT) or a variant where all eighteen cytoplasm-facing lysine residues (listed in Methods) where mutated to arginine (cK0). IMCD3-[pCrys-GPR161^3NG^] cells were treated with SAG for 2h. During the course of the experiment, NG fluorescence was tracked in individual cilia at 10 min capture intervals. Fluorescence data were acquired and analyzed as in panel **(A)**. A comparison of the ciliary levels of GPR161 at t=0 is shown in **Fig. S2B**. Error bars: 95% CI. *N* =10-19 cilia. **C.** IMCD3-[^AP^SSTR3; BirA-ER] cells expressing either WT or cK0 SSTR3 were pulse-labeled by addition of Alexa647-conjugated mSA (mSA647) to the medium for 5 min before addition of sst and were imaged by far red fluorescence immediately after addition of sst (t_0_) and 10 min later (t_10_). The contrast level was adjusted to reveal plasma membrane-localized and internalized ^AP^SSTR3, causing the cilia-localized ^AP^SSTR3 signal to reach saturation. Scale bar 5μm. **D.** Internalized ^AP^SSTR3 foci were counted immediately after addition of sst (t_0_) and 10 min later (t_10_), n = 34 cells.

To rule out that the observed ciliary exit defect of the cK0 mutants was an indirect consequence of defective endocytosis for example because of clogging of the ciliary exit path, we directly assessed endocytosis of SSTR3 and SSTR3cK0. The surface-exposed pool of ^AP^SSTR3 was pulse labeled with fluorescently labeled monovalent streptavidin (mSA) and cells were stimulated with sst. The faint hazy signal corresponding to the plasma membrane pool of SSTR3 disappeared after 10 min in the presence of sst in a similar fashion for both WT and cK0 variants (**Figure 2C**). Concurrently, the WT and cK0 variants appeared in cytoplasmic foci corresponding to endocytic vesicles (**Figure 2C**). Counting cytoplasmic foci revealed that endocytosis proceeded normally irrespective of the ubiquitination competence of SSTR3 (**Figure 2D**). SSTR3 thus behaves similarly to nearly every GPCR tested to date, in that it does not require ubiquitination for its signal-dependent internalization. Indeed, while ubiquitination of plasma membrane proteins is a major driver of internalization in yeast, ubiquitination of signaling receptors is generally dispensable for signal-dependent internalization in mammalian cells (Dores and Trejo, 2019; Piper et al., 2014; Skieterska et al., 2017).

These results strongly support a direct role for GPCR ubiquitination in promoting signal-dependent exit from cilia.

### Ciliary K63 ubiquitin linkages are required for GPCR exit

We next sought to characterize the type of ubiquitin conjugates that are attached to ciliary proteins. The FK1 and FK2 antibodies used in our immunofluorescent studies recognize ubiquitin either as single adduct or in chain but not free ubiquitin (Fujimuro et al., 1994; Emmerich and Cohen, 2015). The availability of antibodies specific for K48 and K63 ubiquitin linkages (Newton et al., 2008) enabled us to determine the types of ubiquitin conjugates that are attached to ciliary GPCRs. While we found no detectable signal for K48-linked ubiquitin chains (UbK48) in cilia, the signal for K63-linked ubiquitin (UbK63) linkages mirrored the ciliary response observed with the FK1 and FK2 antibody in IMCD3-[SSTR3] cells subjected to agonist treatment (**Figure 3A-B and S1C-D**). Given the high specificity and selectivity of the K48 and K63 linkage antibodies, these data strongly suggest that K63Ub chains are added onto SSTR3 inside cilia upon activation.

**Figure 3.**
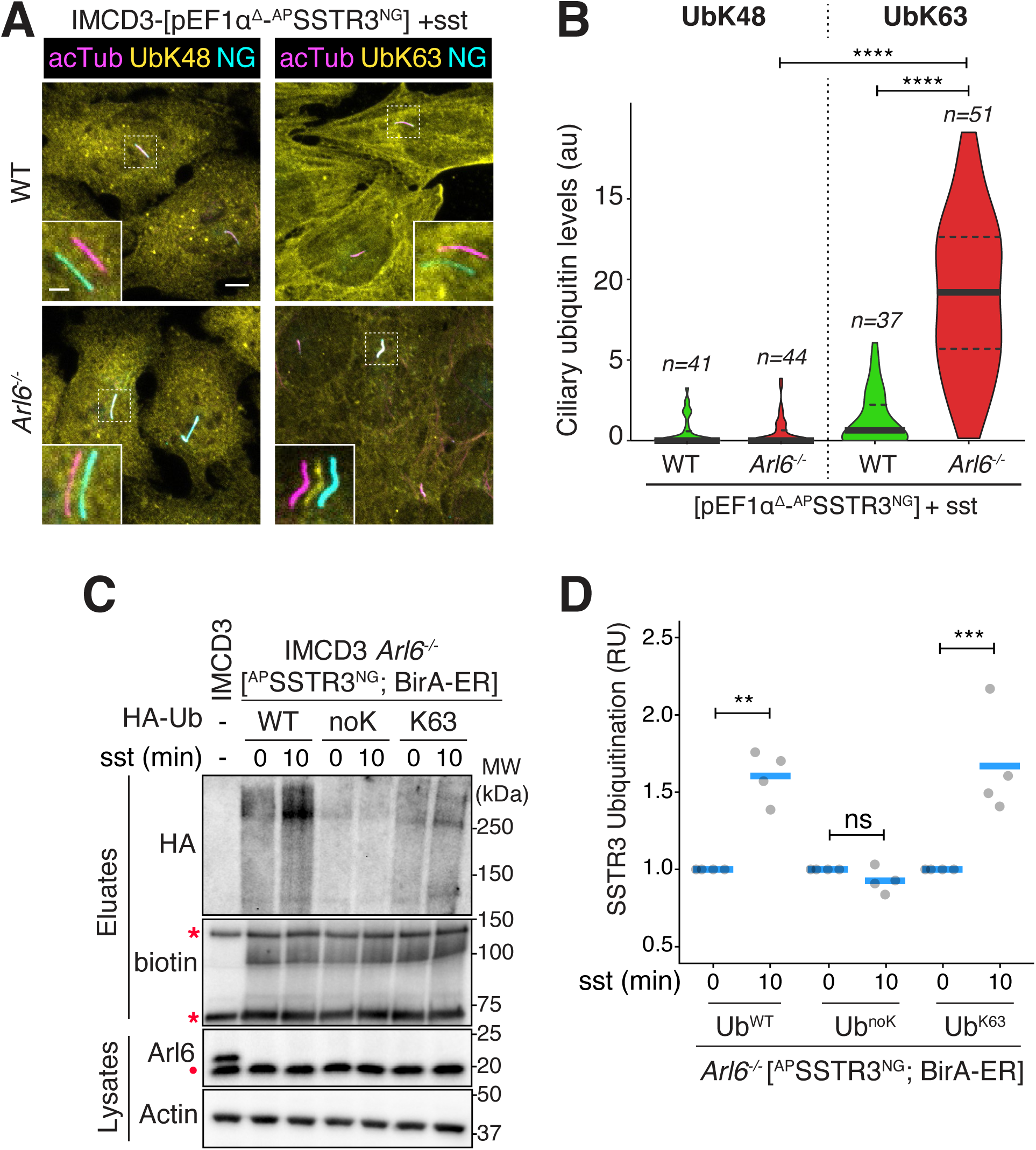
Signal-dependent accumulation of K63-linked ubiquitin chains inside cilia of *Bbs* mutant cells. **A.** IMCD3 cells of the indicated genotypes expressing ^AP^SSTR3^NG^ were treated with somatostatin-14 for 2 h. Cells were fixed and stained for AcTub (magenta) and with antibodies specific for the lysine 63 (UbK63) or lysine 48 (UbK48) Ub chain linkages (yellow). SSTR3^NG^ (cyan) was imaged through the intrinsic fluorescence of NG. Channels are shifted in the insets to facilitate visualization of overlapping ciliary signals. Scale bar 5μm (main panel), 2μm (inset). **B.** The fluorescence intensity of the UbK48 and UbK63 channels in the cilium are represented as violin plots. A 14-fold increase in ciliary Ub abundance is detected with the K63Ub linkage-specific antibody. Asterisks indicate ANOVA significance value. ****, p= <0.0001. No ciliary signal is detected with the K48Ub linkage-specific antibody. **C.** *Arl6^-/-^* IMCD3-[^AP^SSTR3; BirA-ER] cells were transfected with the HA-tagged ubiquitin variants WT, noK0 (all seven acceptor lysine residues mutated to arginine) or K63 (where all lysine residues are mutated to arginine except for K63). Biotin was included in the culture medium and cells were treated with sst for 0 or 10 min. Cells were lysed under denaturing conditions and biotinylated SSTR3 was captured on streptavidin resin. Eluates were probed for HA via immunoblotting and for biotin via streptavidin-HRP. Two major biotinylated proteins endogenous to cells are marked by asterisks. Whole cell lysates were probed for Arl6 and, as a loading control, actin. A non-specific band cross-reacting with the anti-Arl6 antibody is marked with a dot. WT IMCD3 cells were processed in parallel as a control. **D.** Quantitation of SSTR3 ubiquitination. The signals of HA-Ub conjugated to SSTR3 in the streptavidin eluates were measured. The experiment shown in **C** was repeated four times and for each Ub variant, Ub-SSTR3 signals were normalized to the value at t = 0 of sst stimulation and plotted as grey circles. The horizontal blue lines represent mean values. Asterisks indicate ANOVA significance value. *** p *=*< 0.001; ** p=<0.01.

**Figure 4.**
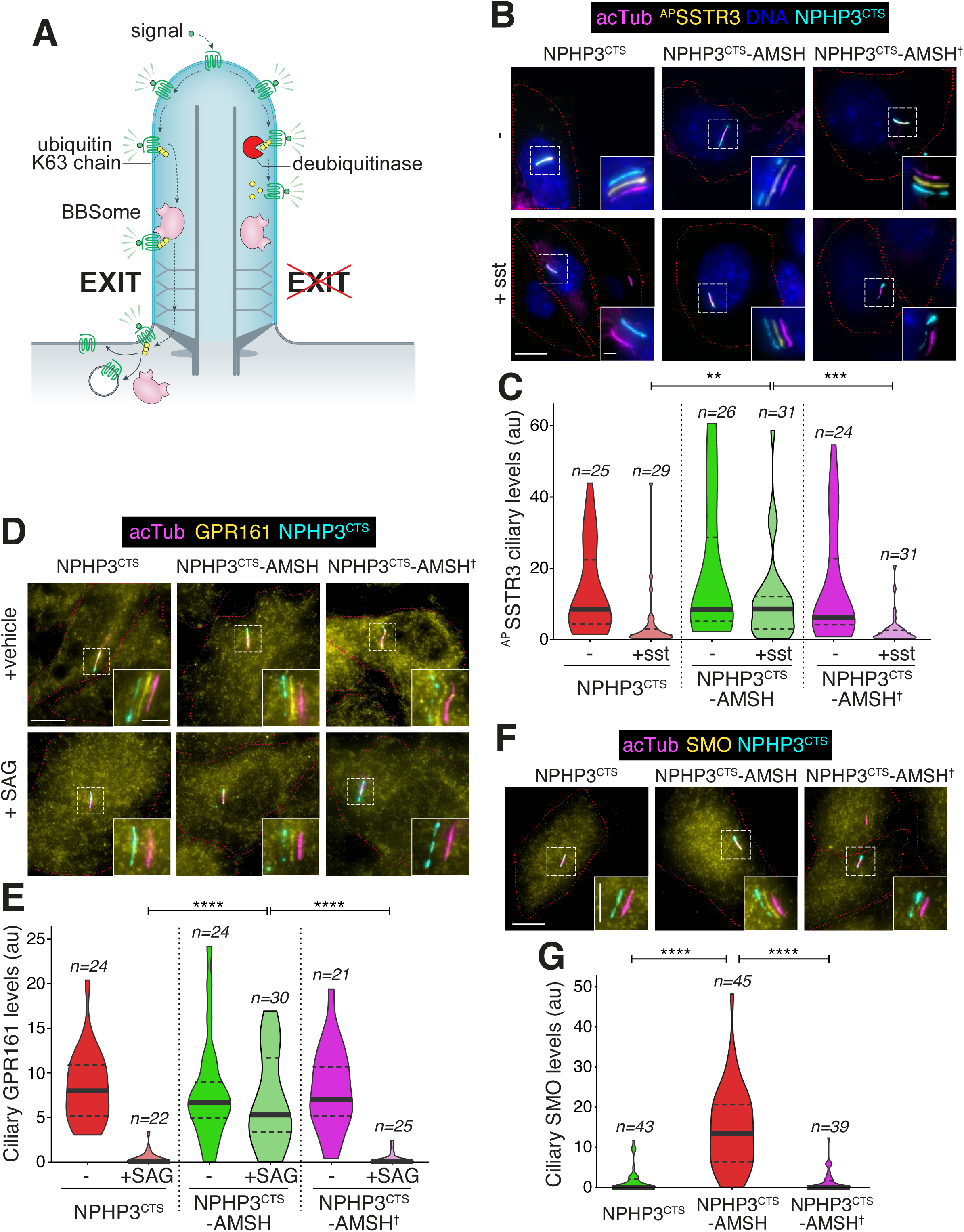
Ciliary K63-linked ubiquitin chains are required for GPCR exit from cili. **A.** Diagram of the working model and the experimental strategy. **B.** IMCD3-[pEF1α^Δ^-^AP^SSTR3; pEF1α-BirA•ER] were transfected with plasmids expressing the ciliary targeting signal (CTS) of NPHP3 fused to GFP only (left panels), to GFP and the catalytic domain of the K63- specific deubiquitinase AMSH (middle panels), or to GFP and the catalytically inactive E280A variant of AMSH catalytic domain (AMSH^†^, right panels). Ciliary ^AP^SSTR3 was pulse-labeled with mSA647 for 5-10 min and cells were then treated with or without sst for 2h, before fixation and staining for acetylated tubulin (acTub, magenta) and DNA (blue). The NPHP3^CTS^ fusions were visualized through the intrinsic fluorescence of GFP (cyan) and ^AP^SSTR3 was visualized via mSA647 (yellow). Channels are shifted in the insets to facilitate visualization of overlapping ciliary signals. Scale bar 5μm (main panel), 2μm (inset). **C.** The fluorescence intensities of ciliary ^mSA647-AP^SSTR3 are represented as violin plots. Asterisks indicate Kruskal-Wallis test significance values. ***, p= <0.001; **, p= <0.01. Addition of sst triggers the exit of SSTR3 from cilia in cells transfected with the controls NPHP3^CTS^ and NPHP3^CTS^-AMSH^†^ but NPHP3^CTS^-AMSH blocks ciliary exit of SSTR3. **D.** RPE1-hTERT (RPE) cells transfected with the indicated constructs were treated with SAG or vehicle (DMSO) for 2h, then fixed and stained for acetylated tubulin (magenta) and GPR161 (yellow). The NPHP3^CTS^ fusions were visualized through the intrinsic fluorescence of GFP (cyan). Channels are shifted in the insets to facilitate visualization of overlapping ciliary signals. Scale bar 5μm (main panel), 2μm (inset). **E.** The fluorescence intensities of ciliary GPR161 are represented as violin plots. Asterisks indicate ANOVA significance value. ****, p= <0.0001. NPHP3^CTS^-AMSH specifically blocks the SAG-dependent exit of GPR161 from cilia. **F.** IMCD3 cells transfected with the indicated constructs were fixed and stained for acetylated tubulin (magenta) and SMO (yellow). The NPHP3^CTS^ fusions were visualized through the intrinsic fluorescence of GFP (cyan). Channels are shifted in the insets to facilitate visualization of overlapping ciliary signals. **G.** The fluorescence intensities of ciliary SMO are represented as violin plots. Asterisks indicate ANOVA significance value. ****, p= <0.0001. NPHP3^CTS^-AMSH specifically blocks the constitutive exit of SMO from cilia and promotes accumulation of SMO in cilia in the absence of pathway activation.

To confirm that K63 of Ub is the main linkage used for the elongation of Ub chains on SSTR3, we transfected variants of ubiquitin with lysines mutated to arginines into *Arl6^-/-^* cells stably expressing SSTR3. Signal-dependent ubiquitination of SSTR3 was abrogated when all seven lysines of Ub were mutated to arginines (**Figure 3C-D**). However, when K63 was the only lysine left intact on Ub, signal-dependent ubiquitination of SSTR3 was restored to the same extent as with WT Ub (**Figure 3C-D**). We conclude that UbK63 chains are assembled onto SSTR3 in response to SSTR3 activation inside cilia.

To determine the functional importance of UbK63 linkages in tagging GPCRs for exit from cilia, we sought to specifically interfere with UbK63 linkages inside cilia (**Figure 4A**). Structural and biochemical studies have demonstrated that the deubiquitinase AMSH possesses a near-absolute specificity for UbK63 linkages and does not cleave other Ub chain linkages or Ub-substrates linkages (Sato et al., 2008; McCullough et al., 2004). AMSH normally function on the surface of late endosomes in concert with the endosomal sorting complex required for transport (ESCRT) protein STAM (McCullough et al., 2006). AMSH was previously fused to the epidermal growth factor receptor EGFR to demonstrate that UbK63 chains assembled on EGFR are dispensable for internalization but required for sorting into lysosomes (Huang et al., 2013). We targeted AMSH to cilia using the well-validated ciliary targeting signal from the ciliopathy protein NPHP3 (Nakata et al., 2012; Wright et al., 2011) and analyzed the exit of three distinct GPCRs: SMO, GPR161 and SSTR3. To alleviate concerns related to the interaction of AMSH with the STAM, we deleted the STAM-interacting domain of AMSH and only expressed the catalytic domain of AMSH. The signal-dependent exit of SSTR3 was assessed by treating IMCD3-[SSTR3] cells with sst for 2h. In cells expressing cilia-AMSH (a fusion of NPHP3[1-200] to GFP and the catalytic domain of AMSH), sst no longer triggered a significant reduction in the ciliary levels of SSTR3 (**Figure 4B-C**). Care was taken to only analyze cells that expressed modest levels of cilia-AMSH as judged by GFP fluorescence (**Figure S3**). In contrast, SSTR3 exit proceeded normally in cells that expressed NPHP3[1-200]-GFP alone or a catalytically dead variant of cilia-AMSH that we named cilia-ASMH^†^ (**Figure 4B-C**). These controls strongly suggest that cilia-AMSH exerts its effects on SSTR3 exit by cleaving UbK63 chains inside cilia, a conclusion borne out by imaging ciliary UbK63 in cells transfected with cilia-AMSH (**Figure S4**). Similarly, the exit of endogenous GPR161 from RPE cell cilia upon activation of the Hedgehog pathway was abolished in cells expressing moderate levels of cilia-AMSH (**Figure 4D-E**). Neither NPHP3[1-200]-GFP nor cilia-AMSH^†^ blocked the signal-dependent exit of GPR161 from cilia (**Figure 4D-E**).

Unlike GPR161 and other ciliary GPCRs such as the Somatostatin receptor 3 (SSTR3) that are removed from cilia by the BBSome only when they become activated, SMO undergoes constitutive BBSome-dependent exit from cilia in the absence of Hedgehog pathway stimulation (Nachury, 2018; Ye et al., 2018; Ocbina and Anderson, 2008; Zhang et al., 2011). Signal-dependent accumulation of SMO in cilia is achieved, at least in part, by suppression of its exit (Milenkovic et al., 2015; Nachury and Mick, 2019; Ye et al., 2018). In cells expressing cilia-AMSH, endogenous SMO spontaneously accumulated inside cilia in the absence of Hedgehog pathway activation (**Figure 4F-G**). Again, neither NPHP3[1-200]-GFP nor cilia-AMSH^†^ influenced the ciliary accumulation of SMO (**Figure 4F-G**).

Together, these results indicate that K63-linked ubiquitin chains inside cilia, presumably built onto each ciliary GPCRs under a specific signaling stimulus, are required for GPCR removal from cilia.

### β-arrestin 2 mediates signal-dependent ubiquitination of ciliary GPCRs

As β-arrestin 2 is required for signal-dependent exit of GPR161 and SSTR3 from cilia (Green et al., 2015; Pal et al., 2016; Ye et al., 2018), we sought to determine the functional relationship between β-arrestin 2 and the BBSome. As previously described (Green et al., 2015), β-arrestin 2 is initially undetectable inside cilia and becomes recruited to cilia within minutes of SSTR3 agonist addition (**Figure 5A-C**). The ciliary β-arrestin 2 signal reaches a plateau after about 10 min and the t_1/2_ for ciliary accumulation is less than 5 min (**Figure 5C**). Meanwhile the t_1/2_ for BBSome recruitment to the tip of cilia was 10 min and the first signs of detectable SSTR3 exit were seen between 10 and 15 min (**Figure 5C**). These kinetics suggested that β-arrestin 2 is first recruited to activated ciliary SSTR3 before the BBSome ferries activated SSTR3 out of cilia. When ARL6 was depleted, the basal levels of ciliary β-arrestin 2 were elevated but the kinetics of β-arrestin 2 accumulation in cilia upon SSTR3 activation were indistinguishable from control-depleted cells (**Figure 5A-B**). These data suggest that a small fraction of SSTR3 that is tonically active fails to exit cilia in ARL6-depleted cells and recruits β-arrestin 2. From these data, we conclude that β-arrestin 2 functions either upstream of, or in parallel with the BBSome in signal-dependent retrieval of GPCRs. Similarly, the kinetics of GPR161 exit and β-arrestin 2 ciliary recruitment upon activation of the Hh pathway indicated that β-arrestin 2 reaches its maximal ciliary level before the first signs of GPR161 exit become evident (**Figure 5D**).

**Figure 5.**
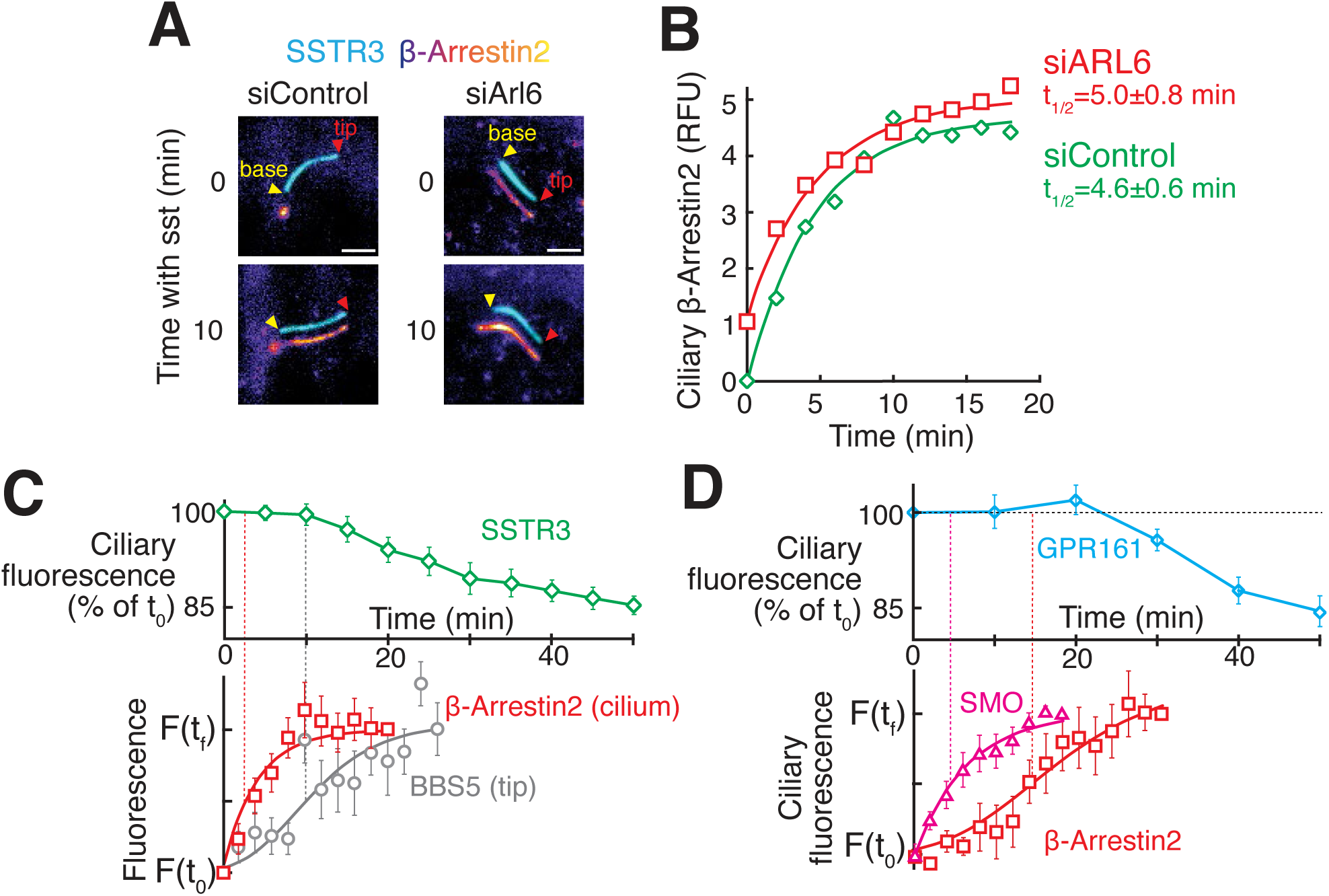
β-arrestin 2 is recruited to cilia upon activation of SSTR3 or GPR161. **A.** β-Arrestin2 is rapidly recruited to cilia upon activation of SSTR3. siRNA-treated IMCD3-[^AP^SSTR3, β-Arrestin2^GFP^] cells were pulse-labeled with mSA647 before addition of sst. Channels are shifted to facilitate visualization of overlapping ciliary signals. As shown previously (Shankar et al., 2010), β-Arrestin2 is at the basal body in unstimulated cells. Scale bar: 2 μm. **B.** Kinetics of β-Arrestin2^GFP^ recruitment to cilia upon sst stimulation in control- and ARL6-depleted IMCD3-[^AP^SSTR3, β-Arrestin2^GFP^] cells. Data were fit to a simple exponential. (n=10 cilia). **C-D.** Kinetics of early trafficking events for retrieval of SSTR3 **(C)** and GPR161 **(D)**. The top panels show removal of stably expressed ^AP^SSTR3^NG^ or ^AP^GPR161^NG3^ following addition of sst **(C)** or SAG **(D)**. The bottom panels show the kinetics of β-Arrestin2^GFP^ entry **(C-D)**, ^AP^Smoothened^NG^ entry **(D)**, and ^NG3^BBS5 tip accumulation **(C)** measured in IMCD3 cells stably expressing the indicated proteins. Fluorescence values are normalized to the initial (t_0_) and final (t_f_) values. Error bars: SEM. (n=6-11 cilia).

We previously proposed that β-arrestin 2 may bridge activated GPCRs to the BBSome (Ye et al., 2018; Nachury, 2018). Yet, we have thus far failed to detect a biochemical interaction between β-arrestin 2 and the BBSome. Furthermore, the continuous distribution patterns of β-arrestin 2 and SSTR3 inside cilia are similar to one another and distinct from the discontinuous pattern of the BBSome (**Figure 5A**), arguing against a direct association of β-arrestin 2 with BBSome/IFT trains inside cilia. We considered the alternative hypothesis that β-arrestin 2 function upstream of the BBSome. Given that β-arrestin 2 functions in the signal-dependent targeting of GPCRs from the plasma membrane to the lysosome by recruiting ubiquitin ligases to activated GPCRs (Henry et al., 2012; Bhandari et al., 2007; Shenoy et al., 2008; Martin et al., 2003), we posited that β-arrestin 2 recognizes activated GPCRs inside cilia to direct their ubiquitination and subsequent selection by the BBSome for removal from cilia. A central prediction of this model is that β-arrestin 2 is required for the ubiquitination of ciliary GPCRs in response to stimulation. To test this prediction, we deleted the β-arrestin 2 gene *Arrb2* in *Arl6* knockout IMCD3 cells (Nager et al., 2017). While the signal-dependent exit of SSTR3 from cilia failed in both *Arl6^-/-^* and *Arl6^-/-^ Arrb2^-/-^* cells (**Figures 1A-B and 6A-B**), ubiquitin staining in cilia and by inference signal-dependent ubiquitination of SSTR3 was only observed in *Arl6^-/-^* cells (**Figures 6A-B**). In *Arl6^-/-^ Arrb2^-/-^* cells, the signal-dependent increase in ciliary ubiquitin signal was no longer observed (**Figures 6A-B**). Similarly, the increase in ciliary ubiquitin signal seen in *Arl6^-/-^* cells treated with the Hedgehog pathway agonist SAG was no longer observed in the absence of β-arrestin 2 (**Figure 6C-D**). These data indicate that, in the absence of BBSome function, ciliary GPCRs become ubiquitinated in response to activation in a β-arrestin 2-dependent manner. We sought to confirm our imaging-based finding with a biochemical analysis of SSTR3 ubiquitination. While SSTR3 ubiquitination was readily detected in *Arl6^-/-^* cells and measurably increased upon stimulation with sst, the deletion of β-arrestin 2 in *Arl6^-/-^* cells drastically reduced SSTR3 ubiquitination levels and no sst-dependent increase of SSTR3 ubiquitination was detected in *Arl6^-/-^/Arrb2^-/-^* cells (**Figures 6A-B**).

**Figure 6.**
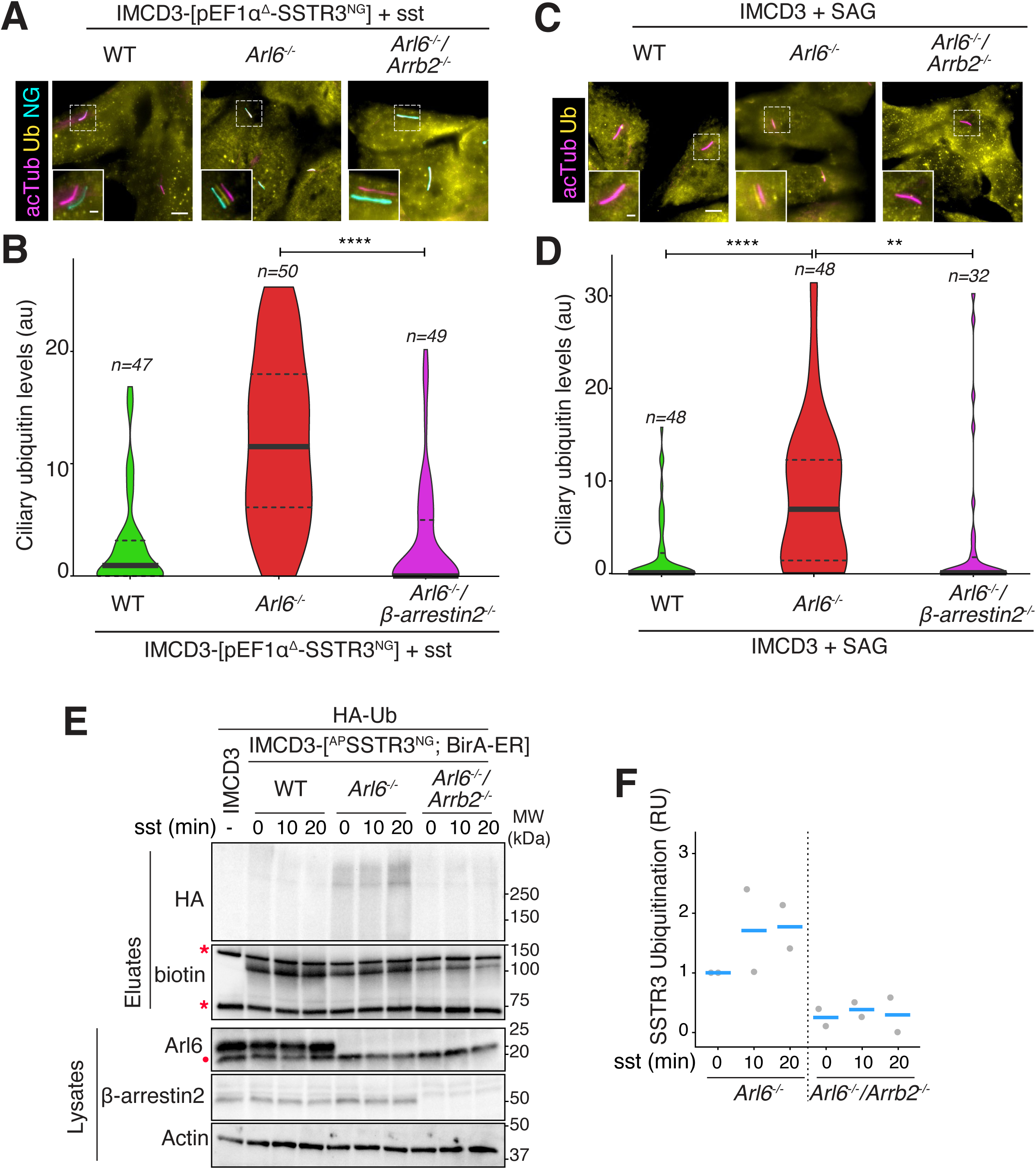
β-arrestin 2 directs the signal-dependent ubiquitination of ciliary GPCRs. **A.** IMCD3-[pEF1α^Δ^-^AP^SSTR3^NG^] cells of the indicated genotypes were treated with sst for 2h before fixation and staining for acetylated tubulin (acTub, magenta) and ubiquitin (Ub, yellow). SSTR3^NG^ was visualized through the intrinsic fluorescence of NG (cyan). Channels are shifted in the insets to facilitate visualization of overlapping ciliary signals. SSTR3 exit is blocked in *Arl6^-/-^* cells regardless of the *β-Arrestin2* genotype but the ciliary Ub signal is only evident when β-Arrestin2 function is intact. Scale bar: 5μm (main panel), 1μm (inset). **B.** Violin plots representing the ciliary levels of Ub under the indicated conditions. Asterisks indicate ANOVA significance value. ****, p= <0.0001. **C.** IMCD3 WT, knockout for *Arl6* only and double knockout for *Arl6* and the β-Arrestin2 gene *Arrb2* were treated with SAG for 2h before fixation and staining for acetylated tubulin (acTub, magenta) and ubiquitin (Ub, yellow). Channels are shifted in the insets to facilitate visualization of overlapping ciliary signals. Scale bar: 5μm (main panel), 1μm (inset). **D.** Violin plots representing the ciliary Ub levels. Asterisks indicate ANOVA significance value. ****, p= <0.0001; **, p= <0.01. The appearance of a Ub signal in cilia in *Arl6^-/-^* cells depends on β-Arrestin2. **E.** IMCD3-[^AP^SSTR3; BirA-ER] cells of the indicated genotypes stably expressing ^AP^SSTR3^NG^ and BirA-ER were transfected with HA-Ub, biotin was added to the medium and cells were treated with sst for indicated times. Biotinylated SSTR3 was captured from cell lysates under denaturing conditions on streptavidin resin. Eluates were probed for HA via immunoblotting and for biotin via streptavidin-HRP. Two major biotinylated proteins endogenous to cells are marked by asterisks. Whole cell lysates were probed for Arl6, β-Arrestin2 to verify genotypes and, as a loading control, actin. A non-specific band cross-reacting with the anti-Arl6 antibody is marked with a dot. WT IMCD3 cells were processed in parallel as a control. **F.** Quantitation of SSTR3 ubiquitination. The signals of HA-Ub conjugated to SSTR3 in the streptavidin eluates were measured. The experiment shown in **E** was repeated twice and for each experiment, Ub-SSTR3 signals were normalized to the value in *Arl6^-/-^* cells at t = 0 of sst stimulation and plotted as grey circles. The horizontal blue lines represent mean values.

Together, these data suggest the following order of action: GPCR activation, β-arrestin 2 engagement, Ub ligase recruitment, assembly of K63Ub chains on the GPCR, and finally selection of ubiquitinated GPCRs for BBSome-mediated retrieval (**Figure 7E**).

**Figure 7.**
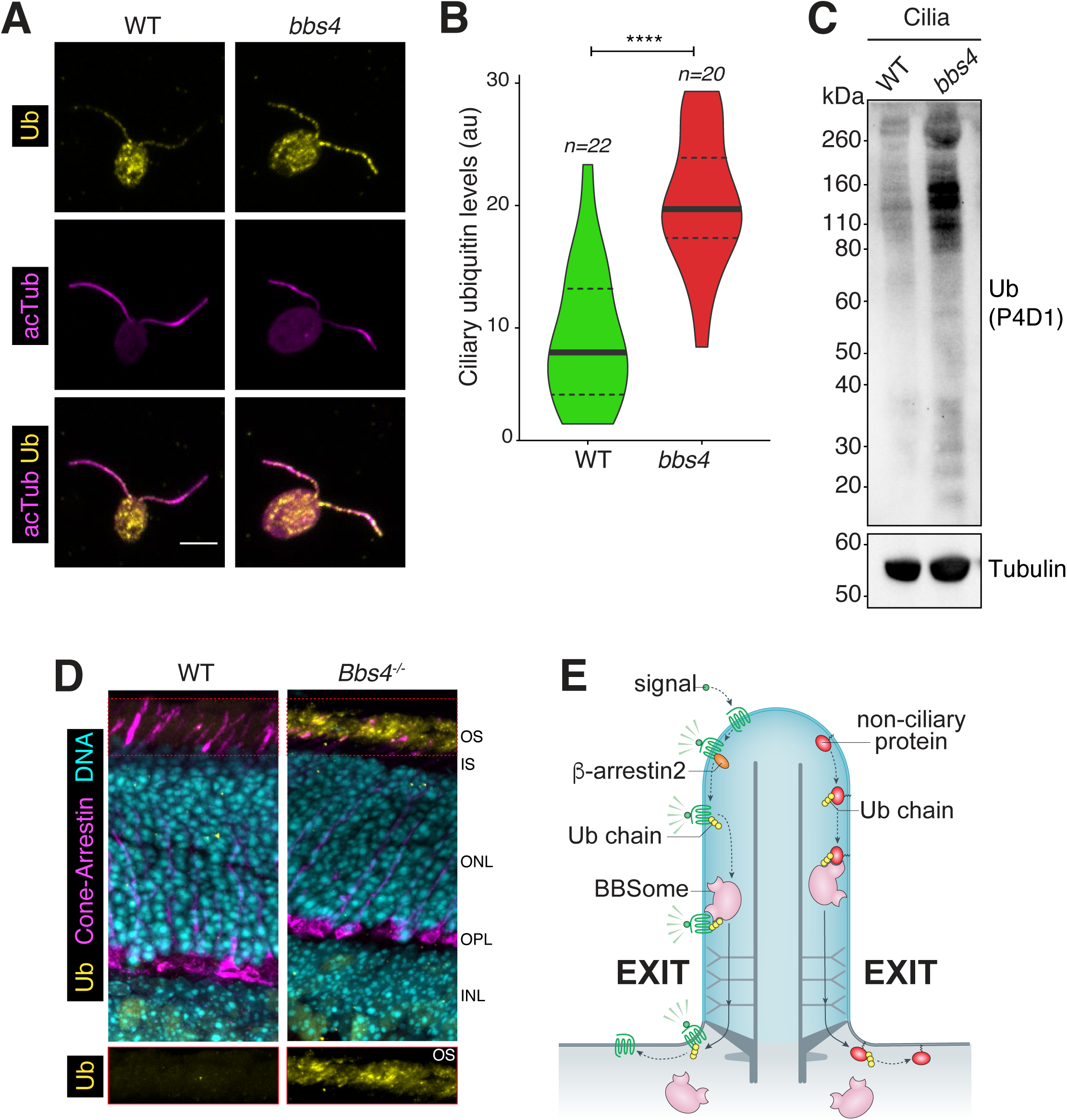
Accumulation of ubiquitinated proteins in cilia of *Chlamydomonas* cells and mammalian photoreceptors. **A.** *Chlamydomonas* cells of the indicated genotypes were fixed and stained for acetylated tubulin (acTub, magenta) and ubiquitin (Ub, yellow). Scale bar: 5 μm. A weak ubiquitin signal detected in WT cells is increased in *bbs4* mutant cells. **B.** Violin plots of the ciliary Ub levels in WT and *bbs4 Chlamydomonas* cells. Asterisks indicate Mann-Whitney test significance values. ****, p= <0.0001. The ciliary levels of Ub are increased more than 2-fold in *bbs4* cells as compared to WT. **C.** Cilia purified from WT and *bbs4* cells via deciliation with dibucaine and differential centrifugation were solubilized, and the protein contents resolved by SDS-PAGE. Immunoblots for Ub (using the P4D1 antibody) and tubulin are shown. A rise in ubiquitin conjugates is observed in *bbs4 Chlamydomonas* cells compared to WT. **D.** Mouse retina were fixed, embedded and sectioned before staining for ubiquitin (Ub, yellow, lower panel), cone arrestin (magenta, a marker of the photoreceptor outer segment layer) and DNA (cyan). OS: outer segment; IS: inner segment; ONL: outer nuclear layer; OPL: outer plexiform layer; INL: inner nuclear layer. Scale bar: 10μm. **E.** Model for the role of ubiquitin chains in selecting cargoes for signal-dependent and constitutive exit via BBSome-mediated transport.

### Constitutively retrieved BBSome cargoes are ubiquitinated prior to exit

Besides removing GPCRs from cilia in a regulated fashion, the BBSome also clears cilia of unwanted proteins that accidentally enter cilia. In the single-cell flagellated organism *Chlamydomonas reinhardtii*, mutations in the BBSome subunit Bbs4 cause a constitutive accumulation of several proteins including Phospholipase D in cilia (Lechtreck et al., 2009). When stained for ubiquitin, *Bbs4 Chlamydomonas* showed a 2-fold enrichment in ciliary signal compared to WT (**Fig. 7A-B**). Isolation of cilia revealed the accumulation of ubiquitin conjugates above 80 kDa in *Bbs4* mutant *Chlamydomonas* cilia (**Fig. 7C**). These results are consistent with the hypothesis that ciliary proteins subject to constitutive retrieval are ubiquitinated prior to their removal from cilia by the BBSome.

In the mammalian retina, proteomics studies have found over 100 proteins accumulating in the outer segment (the equivalent of the cilium) of *Bbs* photoreceptors compared to WT photoreceptors (Datta et al., 2015). When we stained sections of mouse retina for ubiquitin, we found a drastic accumulation of ubiquitin signal in the outer segment of *Bbs4^-/-^* photoreceptors compared to WT photoreceptors (**Fig. 7D**). We note that the nature of ubiquitin conjugates remains to be determined in the case of constitutive cargoes. These results suggest that a variety of proteins that accidentally flow into the photoreceptors outer segments –possibly caught on the tremendous flux of rhodopsin trafficking to the outer segment (Baker et al., 2008)– are recognized as foreign to the cilium, ubiquitinated on site and selected for removal by the BBSome (**Figure 7E**).

## DISCUSSION

### Recognition of K63-linked ubiquitin chains by the ciliary exit machinery

Our results indicate that K63-linked ubiquitin chains tag activated GPCRs and likely other unwanted proteins for BBSome-mediated removal from cilia. Prior findings in trypanosomes suggest that the BBSome may directly recognize ubiquitin (Langousis et al., 2016), although direct biochemical evidence and information regarding ubiquitin chain specificity is still missing. It is also known that the BBSome directly recognizes cytoplasmic determinants in the GPCRs it ferries out of cilia (Jin et al., 2010; Klink et al., 2017; Ye et al., 2018; Seo et al., 2011; Domire et al., 2011). We propose that the summation of weak molecular interactions between the BBSome and UbK63 chains on one hand and the BBSome and GPCRs cytoplasmic loops on the other hand increases the BBSome-cargo interaction to enable sorting of ubiquitinated GPCRs out of cilia. In support of this bivalent recognition of ubiquitinated cargoes by the BBSome, the affinity of the BBSome for cytoplasmic determinant of GPCRs are in the μM range (Klink et al., 2017) and affinities of ubiquitin-binding proteins for ubiquitin typically reside in the sub mM range (Husnjak and Dikic, 2012). The nature of the biochemical entity that directly recognizes ubiquitin, and in particular UbK63 chains remains to be determined: while the BBSome was pulled down on Ub-agarose resin from trypanosome lysates (Langousis et al., 2016), no known Ub-binding domain is found in the BBSome polypeptides. Furthermore, the IFT-A subunit IFT-139 associates with ubiquitinated α-tubulin in ciliary extracts from *Chlamydomonas*, suggesting a possible recognition of ubiquitin by IFT139 (Wang et al., 2019). Again, the absence of known Ub-binding domain in the sequence of IFT139 presents a challenge to this hypothesis.

### Requirements for K63-linked ubiquitination in ciliary exit and sorting to the lysosome

Wang et al. (2019) found that α-tubulin is modified by UbK63 chains and not UbK48 chain upon ciliary disassembly. The ubiquitin ligases that have been associated with ciliary clearance of Patched 1 and of PDGFRαα specifically assemble UbK63 chains (Kim et al., 2015; Yue et al., 2014; Schmid et al., 2018). The suppression of GPR161, SSTR3 and SMO exit by cilia-AMSH (**Figure 4**) demonstrates that UbK63 chains are recognized inside cilia to direct trafficking out of cilia. The milder exit defect observed when acceptor lysines are mutated on ciliary GPCRs (**Figure 2**) than when cilia-AMSH is expressed (**Figure 4**) suggests that K63Ub chains added onto other ciliary proteins besides GPCRs may participate in ciliary exit. A role for ubiquitination of the ciliary transport machinery in exit is in line with the functional importance of ubiquitination of the endolysosomal machinery in GPCR trafficking (Dores and Trejo, 2019). β arrestin ubiquitination promotes internalization of the β_2_-adrenergic receptor (β_2_AR) (Shenoy et al., 2001, 2007; Shenoy and Lefkowitz, 2003), ubiquitination of the ESCRT components Hrs and STAM participates in sorting of the chemokine receptor CXCR4 into the MVB (Malik and Marchese, 2010; Marchese et al., 2003) and ubiquitination of the ESCRT-III recruitment factor ALIX enhances MVB sorting of the protease activated receptor PAR-1 (Dores et al., 2015).

The requirement for UbK63 chains in signal-dependent exit from the cilium contrasts with the more limited role of ubiquitin and UbK63 chains in signal-dependent endocytosis of signaling receptors at the plasma membrane in mammals. Although ubiquitination is a major driver of endocytosis in yeast (Stringer and Piper, 2011; Piper et al., 2014), foundational studies on epidermal growth factor receptor (EGFR) trafficking found that signal-dependent receptor ubiquitination is not essential for internalization because it is redundant with other internalization mechanisms (Goh et al., 2010; Huang et al., 2013; Fortian et al., 2015). Studies of an EGFR-AMSH fusion found that UbK63 chains attached onto EGFR become increasingly necessary as EGFR progresses along the degradative route and a strict requirement for K63-linked ubiquitin chains is observed at the terminal step of EGFR sorting into the lumen of multivesicular body (MVB) (Huang et al., 2013). A strict requirement for receptor ubiquitination in degradative sorting but not in internalization can be generalized to nearly all GPCRs (Skieterska et al., 2017), namely β_2_AR (Shenoy et al., 2001), CXCR4 (Marchese and Benovic, 2001), µ opioid receptor (Hislop et al., 2011), κ opioid receptor (Li et al., 2008), δ opioid receptor (Henry et al., 2011), neurokinin-1 receptor (Cottrell et al., 2006), protease activated receptor 2 (PAR-2) (Jacob et al., 2005) and vasopressin receptor 2 (V2R) (Martin et al., 2003). Similarly, we find that SSTR3 endocytosis proceeds normally in the absence of SSTR3 ubiquitination (**Figure 2C-D**). Studying the importance of ubiquitin K63 in mammalian cells via genetics remains challenging because of the multiple genes encoding Ub but in yeast strains expressing UbK63R as the sole source of Ub, the Gap1 permease is internalized normally and fails to get sorted into the MVB (Lauwers et al., 2009).

The parallels between sorting at the late endosome and ciliary trafficking extend to β arrestin-directed ubiquitination. While β arrestin functions inside cilia to direct the addition of ubiquitin onto activated SSTR3 and GPR161 (**Figures 5 and 6**), β arrestin-mediated ubiquitination selectively affects sorting at the level of endosomes rather that at the plasma membrane for activated CXCR4 (Bhandari et al., 2007), β_2_AR (Shenoy et al., 2008) and V2R (Martin et al., 2003) and β arrestin-mediated ubiquitination of CXCR4 was shown to take place on endosomes (Malik and Marchese, 2010). In this context, the recognition of K63-linked ubiquitin chain inside cilia may befit the endosomal origin of primary cilia (Sorokin, 1962; Westlake et al., 2011; Mitchell, 2017).

The parallels between endosomal and ciliary trafficking lead us to speculate that some ubiquitinated proteins might be accidentally sorted to cilia instead of late endosomes. These proteins would then need to be retrieved from cilia by the BBSome. Thus, the Ub signal detected in photoreceptor outer segment and in *Chlamydomonas* flagella might result from the accidental import of ubiquitinated proteins into cilia rather than *in situ* ubiquitination of unwanted proteins inside cilia.

### Potential coupling between BBSome-mediated exit and degradative sorting

An interesting consideration is the relationship between ciliary exit and endo-lysosomal sorting. It has been proposed that endocytosis of signaling receptors is intimately coupled to their exit from cilia (Pedersen et al., 2016). However, single molecule imaging of GPR161 exiting cilia suggests that activated GPR161 diffuses into the plasma membrane after it exits from the ciliary compartment (Ye et al., 2018). Prior findings that worm mutants for BBSome or the ESCRT machinery both fail to remove ciliary proteins fused to Ub from cilia (Hu et al., 2007; Xu et al., 2015) suggest the provocative possibility that ciliary exit may be tightly coupled to degradative sorting. Nonetheless, the hypothesis that interfering with UbK63 chain formation on ciliary GPCRs indirectly blocks ciliary exit because of a primary defect in endocytosis can be rejected because ubiquitination is not a major determinant of SSTR3 endocytosis and because cilia-AMSH potently and specifically blocks the signal-dependent exit of ciliary GPCRs.

In this context, the contrast between the dramatic increase in ubiquitinated SSTR3 levels in *Arl6^-/-^* cells compared to WT cells even in the absence of stimulation (**Figure 1C-D**) and the modest increase in ciliary Ub levels in *Arl6^-/-^* cells compared to WT cells in the absence of stimulation (**Figure 1A-B**) suggests that a considerable amount of ubiquitinated SSTR3 accumulates outside of cilia in *Arl6^-/-^* cells. One interpretation consistent with a direct coupling between ciliary exit and degradation is that a small fraction of ubiquitinated SSTR3 escapes cilia in *Arl6^-/-^* cells but fails to reach to correct degradative route. In this interpretation, the BBSome would deliver its cargoes to the ESCRT machinery for routing into the lysosomal degradative pathway.

### Ubiquitination and control of the ciliary proteome

A role for ubiquitination in selecting unwanted proteins for removal from cilia may unify the functions of the BBSome in signal-dependent retrieval of GPCRs and the constitutive clearance of proteins that accidentally enter cilia. In both of these cases, unwanted ciliary proteins need to be recognized as ‘non-self’ by a ciliary surveillance machinery. For GPCRs, β arrestin orchestrate the recognition of activated GPCRs as unwanted by directing their ubiquitination. A fascinating question for future investigations is how the more than 100 proteins that accumulate in outer segments of *Bbs* photoreceptors are recognized as ‘non-self’ and tagged with ubiquitin.

## ACKNOWLEDGMENTS

We thank Drs. Kirk Mykytyn, Val Sheffield, Alice Ting, and Mark Scott for the gifts of cDNAs; Kathryn Anderson, Vishva Dixit, and Felice Dunn for the gifts of antibodies; Val Sheffield for the gift of the *Bbs4^-/-^* mice, George B. Witman and the Chlamydomonas Resource Center for the gift of *Chlamydomonas* strains; Yien-Ming Kuo for help with cryosections of mouse retina and core support; Dhivya Kumar and Tyler Picariello for the assistance with *Chlamydomonas* cultures; Nicholas Morante and Jeremy Reiter for providing *Bbs4^-/-^* mouse eyes; Fan Ye for generating some of the cell lines used in the study; Irene Ojeda Naharros for help with single cilia tracking, the Nachury and von Zastrow labs for helpful discussions and the Ogden and Caspary labs for comments on the preprint. This work was funded by NIGMS (R01-GM089933, M.V.N.). This work was made possible, in part, by NEI EY002162 - Core Grant for Vision Research and by the Research to Prevent Blindness Unrestricted Grant (M.V.N.). Authors contributions: SRS conducted all experimental work except for the experiments in Figure 5 which were conducted by ARN. MVN supervised research. SRS and MVN wrote the manuscript.

## MATERIALS AND METHODS

### Cell culture

The mouse IMCD3 cell lines used in the study were generated from a parental IMCD3-FlpIn cell line (gift from P.K. Jackson, Stanford University, Stanford, CA). IMCD3-FlpIn cells were cultured in DMEM/F12 (11330-057; Gibco) supplemented with 10% FBS (100-106; Gemini Bio-products), 100 U/ml penicillin-streptomycin (400-109; Gemini Bio-products), and 2 mM L-glutamine (400-106; Gemini Bio-products). The RPE1-hTERT cell line (ATCC CRL-4000) was cultured in DMEM/F12 supplemented with 10% FBS, 100 U/ml penicillin-streptomycin, 2 mM L-glutamine and 0.26% sodium bicarbonate (25080; Gibco).

Ciliation was induced by serum starvation in media containing 0.2% FBS for 16 to 24 h.

### Plasmid construction and Generation of stable cell lines

Stable isogenic IMCD3 cell lines were generated using the Flp-In system (ThermoFisher Scientific). Low-expression promoters and additional expression cassettes were introduced into pEF5/FRT plasmid as described (Nager et al., 2017; Ye et al., 2018). To reduce expression levels, the EF1α promoter was attenuated by mutating the TATA-box (pEF1α^Δ^) or replaced with a minimal chicken lens δ-crystallin promoter (pCrys).

Coding sequences were amplified from plasmids encoding mouse GPR161 (BC028163; Mammalian Gene Collection [MGC]; GE Healthcare), mouse SSTR3 (gift from Kirk Mykytyn, Ohio State University, Columbus, OH), and human BBS5 (gifts from V. Sheffield, University of Iowa, Iowa City, IA), BirA-ER (gift from Alice Ting). β-arrestin 2 (a gift from Mark Scott) was stably expressed from a pCMV-based plasmid (pEGFP-N). SSTR3 and SMO expression were driven by pEF1α^Δ^, GPR161 expression by pCrys, and BBS5 expression by pEF1α. Coding sequences were fused to GFP, mNeonGreen (NG, (Shaner et al., 2013), or an acceptor peptide for the biotin ligase BirA (AP) (Howarth and Ting, 2008). Multiple rounds of site-directed mutagenesis were performed to generate SSTR3-cK0, a variant with all the cytoplasm-facing lysine residues (K233, K330, K356, K407, and K421) mutated to arginine. GPR161-cK0, a variant with 18 cytoplasm-exposed lysine residues (K83, K84, K93, K167, K247, K250, K269, K296, K358, K362, K455, K469, K473, K481, K486, K497, K504, and K548) mutated to arginine, was gene synthesized (GenScript). Cilia-AMSH was generated by fusing the catalytic domain of mouse AMSH (gift from David Komander; Addgene plasmid #66712; (Michel et al., 2015) with NPHP3(1-200) and GFP to create NPHP3[1-200]-GFP-AMSH[243-424]. A catalytically dead version of AMSH was generated by mutating the water-activating Glu280 residue to Ala (Sato et al., 2008; Huang et al., 2013). pRK5-HA-Ubiquitin WT, K0 (all seven acceptor lysine residues – K6, K11, K27, K29, K33, K48, and K63, are mutated to arginine), and K63 only (all lysines except K63 mutated to arginines)(gift from Ted Dawson; Addgene plasmid numbers – 17608, 17603, and 17606 respectively).

CRISPR-edited *Arl6^-/-^* and *Arl6^-/-^/Arrb2^-/-^* cell lines were described previously (Liew et al., 2014; Nager et al., 2017). The genotypes are *Arl6*, NM_001347244.1:c.- 10_25del; c.3_6del; and *Arrb2*, NM_001271358.1:c.1 12del;c112_113del.

### Transfection

For generation of all stable cell lines except for the ones expressing β-arrestin 2^GFP^, FRT-based plasmids were reverse transfected using XtremeGene9 (Roche) into IMCD3 Flp-In cells along with a plasmid encoding the Flp recombinase (pOG44) and stable transformants were selected by blasticidin resistance (4 µg/ml). The transfection mixture was then added to the cells. pEGFP-N• β-arrestin 2-GFP was transfected into IMCD3 cells using Lipofectamine 2000, and clones were selected using neomycin resistance.

### Antibodies and drug treatments

The following monoclonal antibodies were used for immunofluorescence: anti-acetylated tubulin (mouse; clone 6-11B-1; Sigma-Aldrich; 1:500), anti-ubiquitin clone FK2 was used in all experiments except where indicated (mouse; D058-3; Medical and Biological Laboratories; 1:500), anti-ubiquitin FK1 in Fig. S1A-B (mouse; D071-3; Medical and Biological Laboratories; 1:500), anti-ubiquitin K48 linkage (rabbit; clone Apu2; 05-1307; Millipore-Genentech; 1:500), anti-ubiquitin K63 linkage (human; clone Apu3; a gift from Genentech; 1:500). The following monoclonal antibodies were used for immunoblotting: anti-ubiquitin (mouse; clone P4D1; 3936; Cell Signaling; 1:1000), anti-α tubulin (mouse; clone DM1A; MS-581-P1ABX; Thermo Scientific;1:1000), anti-HA (mouse; clone 16B12; 901501; Biolegend;1:500), streptavidin-HRP (Pierce-21140;Thermo Scientific; 1:10000). The following polyclonal antibodies were used for immunofluorescence: anti-GPR161 (rabbit; 13398-1-AP; Proteintech; 1:100), anti-Smoothened (rabbit; a gift from Kathryn Anderson, Memorial Sloan Kettering Cancer Center, New York, NY; 1:500), anti-cone-arrestin (rabbit; AB15282; Millipore; 1:500). Biotinylated SSTR3 and GPR161 were detected using Alexa Fluor 647–labeled monovalent streptavidin (mSA647) (Ye et al., 2018). The following reagents were used at the indicated concentrations: 200 nM SAG and 10 µM somatostatin 14. Somatostatin 14 stocks were made in DMEM/F12 media, HEPES, no phenol red (11039-021, GIBCO). SAG was dissolved in DMSO (276855, Sigma-Aldrich).

### Imaging and microscopy

Cells were imaged either on a DeltaVision system (Applied Precision) equipped with a PlanApo 60x/1.40 objective lens (Olympus), CoolSNAP HQ2 camera (Photometrics), and solid-state illumination module (InsightSSI), or on a confocal LSM 700 (Zeiss) microscope equipped with 40x Plan-Apochromat 1.3 DIC oil objective. Z stacks were acquired at 0.5 μm interval (LSM 700, Figs. 1, 3, 7, and S1) or 0.2 μm interval (DeltaVision, Figs. 4, 6, S3, and S4).

For fixed imaging, 60,000 cells were seeded on acid-washed coverglass (12 mm #1.5; 12-545-81; Thermo Fisher Scientific). Cells were grown for 24 h and then starved for 16-24 h in 0.2% FBS media before experimental treatment. After treatment, cells were fixed with 4% paraformaldehyde in PBS for 15 min at 37°C and extracted in -20°C methanol for 5 min. Cells were then permeabilized in IF buffer (PBS supplemented with 0.1% Triton X-100, 5% normal donkey serum (017-000-121; Jackson ImmunoResearch Laboratories), and 3% bovine serum albumin (BP1605-100; Thermo Fisher Scientific)), incubated at room temperature for 1h with primary antibodies diluted in IF buffer, washed three times in IF buffer, incubated with secondary antibodies (Jackson ImmunoResearch Laboratories) diluted in IF buffer for 30 min, and washed three times with IF buffer. DNA was stained with Hoechst 33258 (H1398; Molecular Probes), cells were washed twice more with PBS, and coverglass mounted on slides using fluoromount-G (17984-25; Electron Microscopy Sciences).

For live-cell imaging, 300,000 cells were seeded on acid-washed 25mm cover glass (Electron Microscopy Sciences). After 24 h of growth, cells were serum starved for 16 h and transferred to the DeltaVision stage for imaging at 37°C within an environmental chamber that encloses the microscope and the stage. Cells were imaged in DMEM/F12 media, HEPES, no phenol red (11039-021, GIBCO). To measure SSTR3 exit, cells expressing ^AP^SSTR3 were first washed three times with PBS and then pulse-labeled with 2µg/ml mSA647 for 5 min at 37°C. Cells were then imaged after addition of sst (for SSTR3) or SAG (for GPR161) for 2 h with an acquisition every 10 min. Each acquisition consisted of a 3-plane Z-stack with each plane separated by 0.5 µm. Transmittance was set to 5% and exposure time to 500 ms.

WT or *Bbs4^-/-^* mice (Mykytyn et al., 2004) eyes were enucleated at P21. Whole eyes were fixed in 4% paraformaldehyde/PBS overnight at 4°C and then washed three times with PBS. The eyes were infiltrated in 30% sucrose/PBS at 4 °C overnight. Eyes were placed in OCT compound embedding medium (4583; Tissue-Tek), frozen on dry ice, and stored at −80 °C. 14 µm sections were cut on a cryostat (CM 1850; Leica). Sections were processed for immunofluorescence as follows. Sections were blocked in blocking solution (50 mM Tris pH7.4, 100 mM NaCl, 0.1% Triton-X 100, 3% normal donkey serum, and 0.1% BSA) for 1 h at room temperature, then incubated with primary antibodies overnight at 4°C, followed by three washes in wash solution (50 mM Tris pH7.4, 100 mM NaCl, and 0.1% TX100). After rinsing in wash solution, sections were incubated with secondary antibodies for 1 h at room temperature. DNA was counterstained using Hoechst 33258 dye, sections washed twice more with PBS, and mounted on slides using Fluoromount-G. Sections were imaged on Confocal LSM 700 (Zeiss) microscope.

### Image analysis

Files were imported from the Deltavision or LSM700 workstations into ImageJ/ Fiji (National Institutes of Health) for analysis. For the quantification of ciliary ubiquitin signal in fixed cells, maximum intensity projections were used. The ciliary ubiquitin intensity was measured using the following equation:

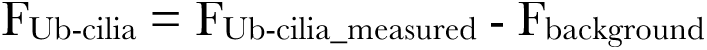

F_Ub-cilia_measured_ is the total ciliary ubiquitin fluorescence detected, F_background_ is the background ubiquitin fluorescence measured in the adjacent area. For all measurements, the fluorescence integrated density was used. F_Ub-cilia_ were plotted as violin plots using the PlotsOfData web tool (https://huygens.science.uva.nl/PlotsOfData/). Each violin represents the distribution of data, which includes all the data points. Median and interquartile range is marked by solid and dotted lines, respectively. No gamma adjustment was applied during figure preparation; all the representative micrographs for respective panels are displayed within the same dynamic range. For some representative micrographs, the most in-focus plane was used rather than the maximum intensity projection (Fig.3, and Fig. 6, panel A and C).

To measure SSTR3 exit, the integrated density of Alexa Fluor 647 fluorescence in the maximum intensity projection was measured for each time point. The cilia-adjacent fluorescence was subtracted as the background, and a mathematical photobleaching correction was applied: F_cilia_ = (F_mSA647_measured_/F_mSA647_1_) + ((1 - e^-λ^) * (n - 1)), where λ is the photobleaching decay constant, n is the number of images taken, F_mSA647_measured_ is the integrated mSA647 fluorescence measured for image n, F_mSA657_1_ is the measurement for the first time point, and F_cilia_ is the reported fluorescence. In this equation, F_cilia_ is reported in relative fluorescence units (RFUs). ^AP^GPR161^3NG^ exit was followed similarly with the difference that NG fluorescence intensity was measured. For SSTR3 as well as GPR161, photobleaching corrected data (F_cilia_) for each condition were linearly fitted (F_cilia_ =m * time + c) and plotted in Excel.

Endocytosed SSTR3 foci were revealed by contrast enhancement and counted with the ImageJ particle analysis tool.

### *Chlamydomonas* culture, isolation of flagella and immunofluorescence

*Chlamydomonas* WT-g1 (nit1, agg1, mt+) [gift from George B. Witman, University of Massachusetts Medical School, Worcester, Massachusetts]and *Bbs4*-CC-4377 ptx5-1/bbs4-1:: NIT1 agg1 mt+ (Chlamydomonas Resource Center) strains were cultured asynchronously under light in TAP media (Gorman and Levine, 1965) for 72 h. Flagellar fractions were prepared from two liters of cultures of WT or *Bbs4* cells as described (Craige et al., 2013). Briefly, cells were harvested by centrifugation for 5 min at 2,000 rpm (1,100 x *g*) at room temperature. Cells were resuspended in 10 mM HEPES, pH 7.4 and centrifuged for 5 min at 2,000 rpm (1,100 x *g*). Next, the cell pellet was gently resuspended in ice-cold HMDS (10 mM HEPES, pH 7.4, 5 mM MgSO_4_, 1 mM DTT, and 4% (w/v) sucrose). Deflagellattion was achieved by addition of 5 mM dibucaine to each tube of cells and by pipetting up and down ∼10 times using a 10-ml plastic serological pipette. The suspension was spun for 5 min at 1,800 x *g* at 4°C. The supernatant containing flagella was underlaid with 9 ml ice-cold HMDS-25% sucrose (10 mM HEPES, pH 7.4, 5 mM MgSO_4_,1 mM DTT, and 25% (w/v) sucrose), followed by a centrifugation step for 10 min at 2,800 rpm (2,400 x *g*), 4°C. The supernatant above the sucrose interface was collected using a 25-ml serological pipette and flagella were pelleted by centrifugation for 20 min at 30,000 x *g* (16,000 rpm), 4°C. The flagellar pellet was resuspended in 100 µl HMDS buffer. Protein concentrations were measured by Bradford and 25 ug were resolved by SDS-PAGE for immunoblotting.

*Chlamydomonas* immunofluorescence was performed as follows. Cells were fixed with 4 % paraformaldehyde in MT buffer* (30 mM HEPES, pH7, 5 mM EGTA, 5 mM MgSO4, and 25mM KCl) for 20 minutes in suspension. The cells were centrifuged at 1000 rpm for 5 minutes, resuspended in 100 µl of fixative and transferred onto slides coated with 1 mg/ml poly-L-lysine. After 5 minutes, the unadhered cells were washed off by rinsing with PBS. Cells were permeabilized in 0.5% Triton-X 100 for 20 minutes followed by blocking for 1 h in blocking buffer (3% fish skin gelatin, 1% BSA, 0.1% Tween20 in PBS) at room temperature. Cells were incubated at 4°C overnight with primary antibodies diluted in blocking buffer, washed five times in PBS and incubated with secondary antibodies for 2 h at room temperature. After five washes in PBS, coverglasses were mounting on slides using Fluoromount-G. Cells were imaged on the LSM700 confocal microscope.

### Biochemical analysis of SSTR3 ubiquitination

IMCD3 cells stably expressing pEF1α^Δ-A^PSSTR3^NG^ and BirA-ER were transiently transfected with HA-tagged ubiquitin plasmids. Cells were reverse transfected using XtremeGene9 and plated into 15 cm plates. Cells were moved to medium containing 0.2% serum and supplemented with 10 µM biotin 24 h after transfection to promote ciliation and maximize biotinylation of SSTR3. 18h after the medium change, sst was added for 0, 10 or 20 min. The assay was stopped by washing cells twice with ice-cold 1xPBS and scraping cells off on ice into 1ml of ice-cold RIPA buffer (50 mM Tris/HCl pH 8, 150 mM NaCl, 1% Nonidet P-40, 0.5% sodium deoxycholate, 0.1% SDS, 100 mM iodoacetamide, 10mM EDTA) supplemented with protease inhibitors (1 mM AEBSF, 0.8 mM Aprotinin, 15 mM E-64, 10 mg/mL Bestatin, 10 mg/mL Pepstatin A and 10 mg/mL Leupeptin). After gently rocking at 4°C for 5 minutes, lysates were clarified by spinning in a tabletop Eppendorf centrifuge at 15,000 rpm and 4°C for 20 minutes. Each clarified lysate was incubated with 50µl of streptavidin Sepharose beads (GE Healthcare) for 1 h at 4°C to capture biotinylated SSTR3. Beads were washed four times with 500µl of RIPA buffer and bound proteins were eluted by heating the beads in 1X LDS sample loading buffer containing 2 mM biotin at 70°C for 10 minutes. After SDS-PAGE and transfer, PVDF membranes were probed with anti-HA antibody and streptavidin-HRP.

Signals were acquired with a ChemiDoc imager (Bio-Rad) and analyzed with ImageLab 6.0.1 (Bio-Rad). The integrated densities of the SSTR3 bands in the streptavidin-HRP blot were normalized to the integrated densities of the endogenous biotinylated proteins to correct for variations in the recovery of biotinylated proteins on streptavidin Sepharose. The resulting value is SSTR3_bio_. Second, the integrated densities of ubiquitinated SSTR3 were collected by measuring the area from 100 kDa to the top of the gel on the HA blot. The values were background-corrected by subtracting the value of the equivalent area in the IMCD3 control lane. The resulting value is Ub-SSTR3_raw_. The relative amount of SSTR3 that is ubiquitinated, Ub- SSTR3_frac_ = Ub-SSTR3_raw_/SSTR3_bio_ was plotted on the graphs using PlotsOfData (Postma and Goedhart, 2019).

---

**Figure S1.**
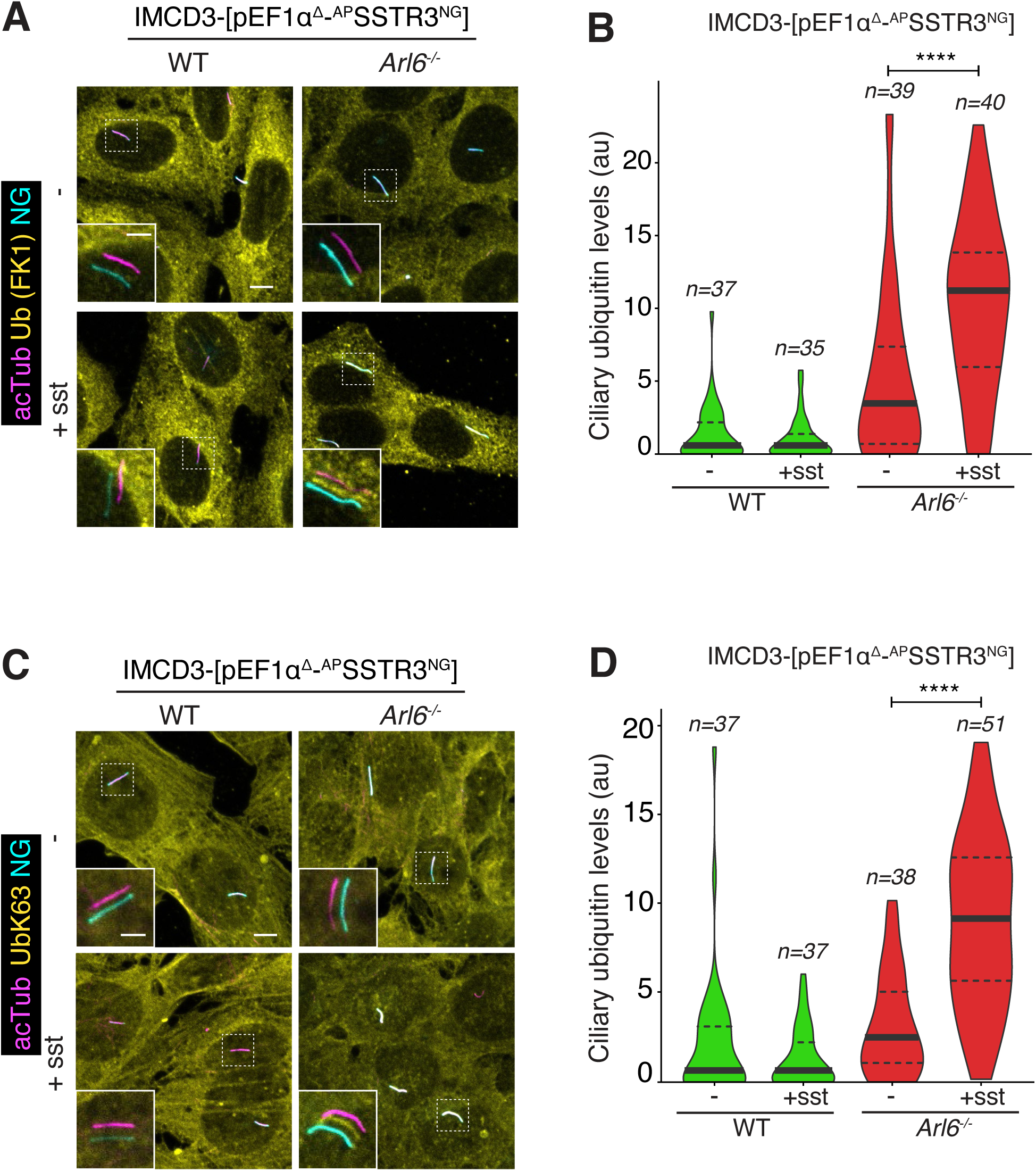
Signal-dependent ubiquitination of ciliary GPCRs. **A.** IMCD3 cells of the indicated genotypes expressing ^AP^SSTR3^NG^ were treated with somatostatin (sst) for 2 h. Cells were fixed and stained for ubiquitin with the FK1 monoclonal antibody (Ub (FK1), yellow) and acetylated tubulin (acTub, magenta). ^AP^SSTR3^NG^ (cyan) was imaged through the intrinsic fluorescence of NG. Channels are shifted in the insets to facilitate visualization of overlapping ciliary signals. Scale bar: 5μm (main panel), 2μm (inset). **B.** Violin plots of the fluorescence intensity of the Ub channel in the cilium in each condition are shown. A 3-fold increase in ciliary Ub signal is observed upon addition of sst to SSTR3-expressing cells. Asterisks indicate ANOVA significance value. ****, p= <0.0001. **C.** IMCD3 cells of the indicated genotypes expressing ^AP^SSTR3^NG^ were treated with sst for 2 h. Cells were fixed and stained for AcTub (magenta) and UbK63 (yellow). ^AP^SSTR3^NG^ (cyan) was imaged through the intrinsic fluorescence of NG. Channels are shifted in the insets to facilitate visualization of overlapping ciliary signals. Scale bar 5μm (main panel), 2μm (inset). **D.** The fluorescence intensity of the UbK63 channel in the cilium is represented as violin plots. Asterisks indicate ANOVA significance value. ****, p= <0.0001.

**Figure S2.**
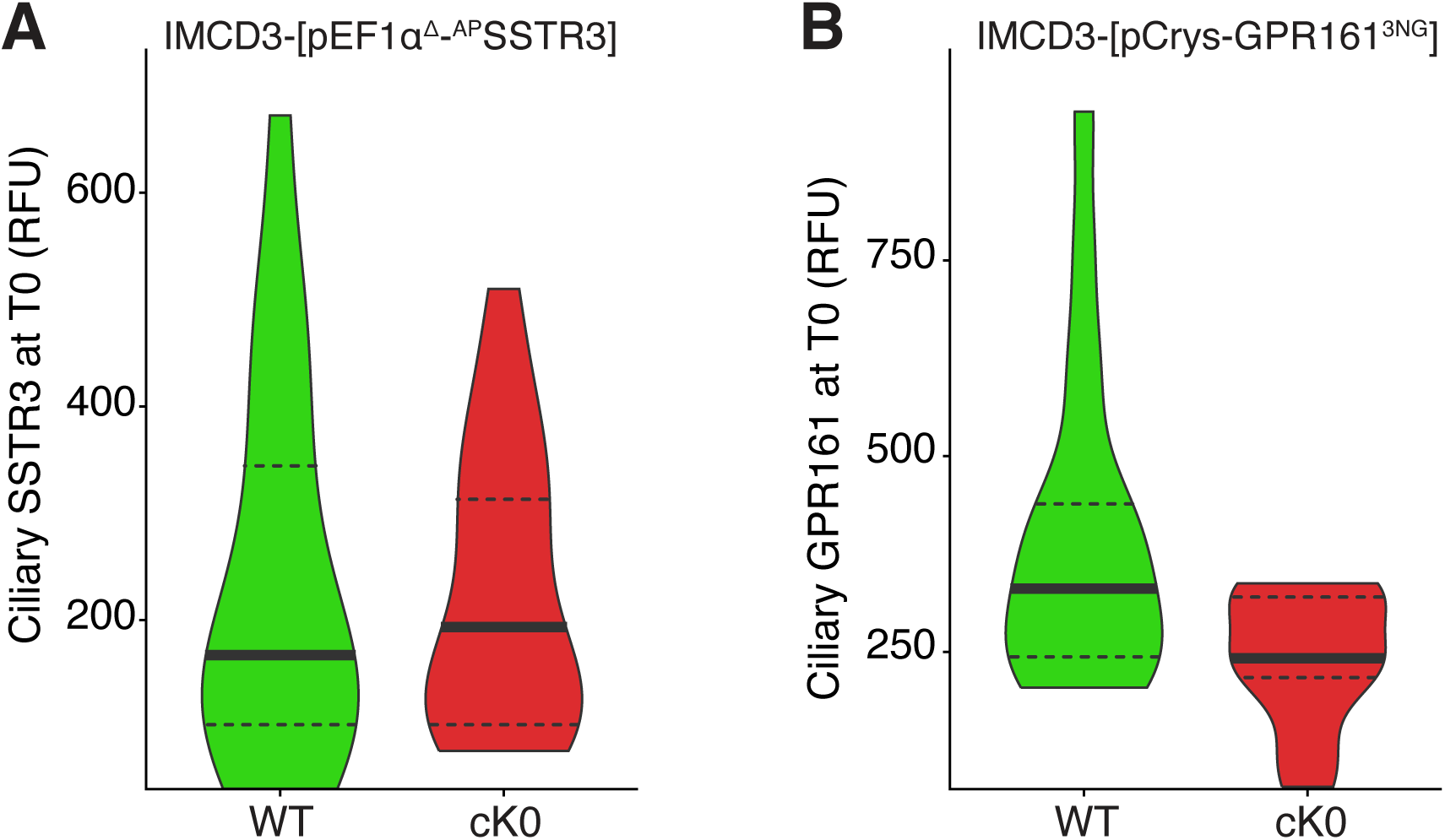
Fluorescence intensities of WT and cK0 GPCR variants at t=0. **A.** Ciliary ^AP^SSTR3 was pulse-labeled by addition of Alexa647-conjugated mSA (mSA647) to the medium for 5-10 min before addition of sst. Violin plots of the ciliary fluorescence intensities for WT and cK0 SSTR3 imaged in the far-red channel at t=0 are shown. **B.** Violin plots of the fluorescence intensities of NG fluorescence corresponding to WT and cK0 GPR161^NG^ at t=0 are shown.

**Figure S3.**
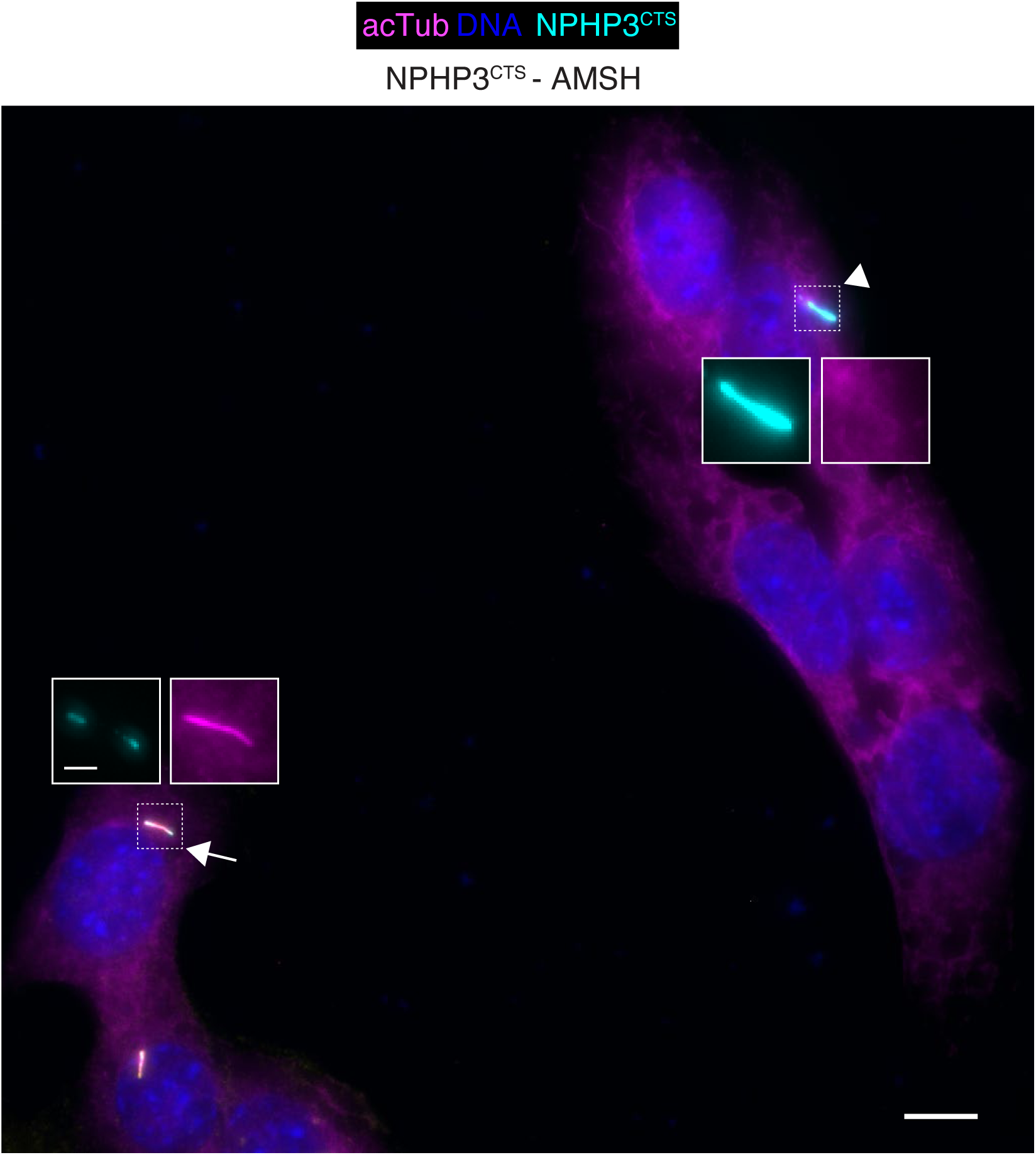
Expression levels of ciliary AMSH. IMCD3-[pEF1α^Δ-AP^SSTR3; pEF1α-BirA•ER] cells were transfected with the plasmid expressing NPHP3^CTS^-AMSH as in Figure 4, before fixation and staining for acetylated tubulin (acTub, magenta) and DNA (blue). The NPHP3^CTS^ fusions were visualized through the intrinsic fluorescence of GFP (cyan) A representative micrograph is shown. A cell expressing modest levels of cilia-AMSH is indicated by an arrow and possesses normal ciliary staining of acetylated tubulin. A cell expressing high levels of cilia-AMSH indicated by an arrowhead displays abnormal ciliary levels of acetylated tubulin. Cells with high ciliary levels of NPHP3^CTS^-AMSH were excluded from the analysis and only cells with modest ciliary levels of NPHP3^CTS^-AMSH were included in the experiment. Scale bar 5µm (main panel), 1µm (inset).

**Figure S4.**
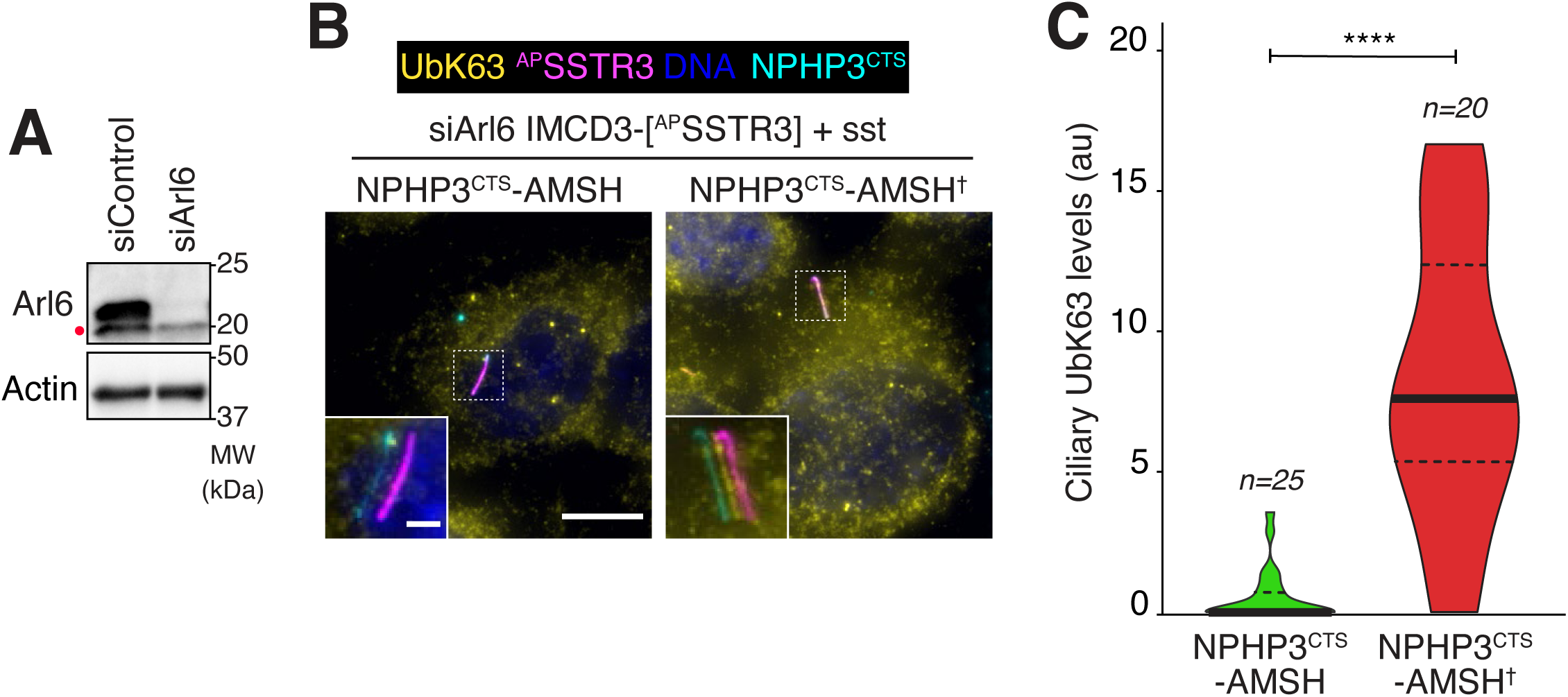
Cilia-targeted AMSH removes UbK63 chains from ciliary substrates. **A.** IMCD3-[pEF1^αΔ-AP^SSTR3; pEF1α-BirA•ER] cells were transfected with control or Arl6 siRNAs. Cell lysates were immunoblotted for Arl6 or actin. A non-specific band cross-reacting with the anti-Arl6 antibody is marked with a dot. **B.** IMCD3-[^AP^SSTR3; BirA•ER] cells were transfected with siRNA targeting Arl6 and plasmids expressing NPHP3^CTS^-AMSH or NPHP3^CTS^-AMSH†. Surface-exposed ^AP^SSTR3 was pulse-labeled with mSA647 for 5-10 min and cells were then treated with sst for 2h before fixation and staining for K63-linked ubiquitin chains (UbK63, yellow). The NPHP3^CTS^-AMSH fusions were visualized through the intrinsic fluorescence of GFP (cyan), ^AP^SSTR3 was visualized via mSA647 (magenta) and DNA is blue. Channels are shifted in the insets to facilitate visualization of overlapping ciliary signals. Scale bar 5μm (main panel), 2μm (inset). **C.** The fluorescence intensities of the UbK63 signal inside cilia are represented as violin plots. Asterisks indicate Mann Whitney test significance value. ****, p= <0.0001.

